# Epigenetic reprogramming shapes the cellular landscape of schwannoma

**DOI:** 10.1101/2022.12.23.521842

**Authors:** S. John Liu, Tim Casey-Clyde, Nam Woo Cho, Jason Swinderman, Melike Pekmezci, Mark C. Dougherty, Kyla Foster, William C. Chen, Javier E. Villanueva-Meyer, Danielle L. Swaney, Harish N. Vasudevan, Abrar Choudhury, Jonathan D. Breshears, Ursula E. Lang, Charlotte D Eaton, Kamir J. Hiam-Galvez, Erica Stevenson, Kuei-Ho Chen, Brian V. Lien, David Wu, Steve E. Braunstein, Penny K. Sneed, Stephen T. Magill, Daniel Lim, Michael W. McDermott, Mitchel S. Berger, Arie Perry, Nevan J. Krogan, Marlon Hansen, Matthew H. Spitzer, Luke Gilbert, Philip V. Theodosopoulos, David R. Raleigh

**Affiliations:** Department of Radiation Oncology, University of California San Francisco, San Francisco, CA 94143, USA; Department of Neurological Surgery, University of California San Francisco, San Francisco, CA 94143, USA; Department of Pathology, University of California San Francisco, San Francisco, CA 94143, USA; Parker Institute for Cancer Immunotherapy, Chan Zuckerberg Biohub, and Departments of Otolaryngology, and Microbiology and Immunology, University of California San Francisco, San Francisco, CA 94115, USA; Department of Urology, University of California San Francisco, San Francisco, CA 94143, USA; Departments of Otolaryngology and Neurosurgery, University of Iowa, Iowa City, IA 52242, USA; Department of Radiology and Biomedical Imaging, University of California San Francisco, USA; J. David Gladstone Institutes, California Institute for Quantitative Biosciences, Department of Cellular and Molecular Pharmacology, University of California San Francisco, San Francisco, CA 94158, USA; Department of Dermatology, University of California San Francisco, San Francisco, CA 94115, USA; Department of Neurological Surgery, Northwestern University, Chicago, IL 60611, USA; Baptist Health Miami Neuroscience Institute, Miami, FL 33176, USA

**Author notes:** Equal contribution (Tim Casey-Clyde, Nam Woo Cho).

## Abstract

Cell state evolution underlies tumor development and response to therapy^1^, but mechanisms specifying cancer cell states and intratumor heterogeneity are incompletely understood. Schwannomas are the most common tumors of the peripheral nervous system and are treated with surgery and ionizing radiation^2–5^. Schwannomas can oscillate in size for many years after radiotherapy^6,7^, suggesting treatment may reprogram schwannoma cells or the tumor microenvironment. Here we show epigenetic reprogramming shapes the cellular landscape of schwannomas. We find schwannomas are comprised of 2 molecular groups distinguished by reactivation of neural crest development pathways or misactivation of nerve injury mechanisms that specify cancer cell states and the architecture of the tumor immune microenvironment. Schwannoma molecular groups can arise independently, but ionizing radiation is sufficient for epigenetic reprogramming of neural crest to immune-enriched schwannoma by remodeling chromatin accessibility, gene expression, and metabolism to drive schwannoma cell state evolution and immune cell infiltration. To define functional genomic mechanisms underlying epigenetic reprograming of schwannomas, we develop a technique for simultaneous interrogation of chromatin accessibility and gene expression coupled with genetic and therapeutic perturbations in single-nuclei. Our results elucidate a framework for understanding epigenetic drivers of cancer evolution and establish a paradigm of epigenetic reprograming of cancer in response to radiotherapy.

## Main text

Cancer is a heterogeneous disease and evolution of cell states and cell types in the tumor microenvironment can influence response to treatment^8–12^. Peripheral nervous system Schwann cells develop from the neural crest^2^, a multipotent embryonic cell population characterized by remarkable molecular and functional diversity^13^. Schwannoma tumors have a low burden of somatic mutations, and schwannoma mutations do not change after treatment with ionizing radiation^14,15^. Unique among human cancers, schwannomas that are treated with radiotherapy undergo long-term oscillations in tumor size^6,7^. In clinical practice, symptomatic schwannoma oscillations are treated with immunosuppressive corticosteroids or surgical decompression, and preoperative schwannoma growth is associated with immune cell infiltration^16,17^. Here we test the hypothesis that epigenetic mechanisms shape the immune microenvironment during schwannoma tumorigenesis and response to radiotherapy. To do so, we interrogate human schwannomas, primary patient-derived schwannoma cells, and schwannoma cell lines using bulk and single-cell bioinformatic, functional genomic, proteomic, metabolomic, and mechanistic approaches. We find schwannomas are comprised of 2 molecular groups that can arise *de novo* (Fig. 1) but undergo epigenetic reprogramming in response to radiotherapy (Fig. 2). Genome-wide CRISPR interference (CRISPRi) screening identifies epigenetic regulators driving schwannoma cell reprogramming and immune cell infiltration in response to ionizing radiation (Fig. 3), and a new technique integrating single-nuclei ATAC, RNA, and CRISPRi perturbation sequencing elucidates concordant chromatin accessibility, transcription factors, and gene expression programs underlying schwannoma cell state evolution that are conserved in human tumors (Fig. 4). In sum, these data shed light on the molecular landscape of schwannomas and reveal epigenetic mechanisms underlying intratumor heterogeneity in response to radiotherapy (Fig. 5).

**Fig. 1.**
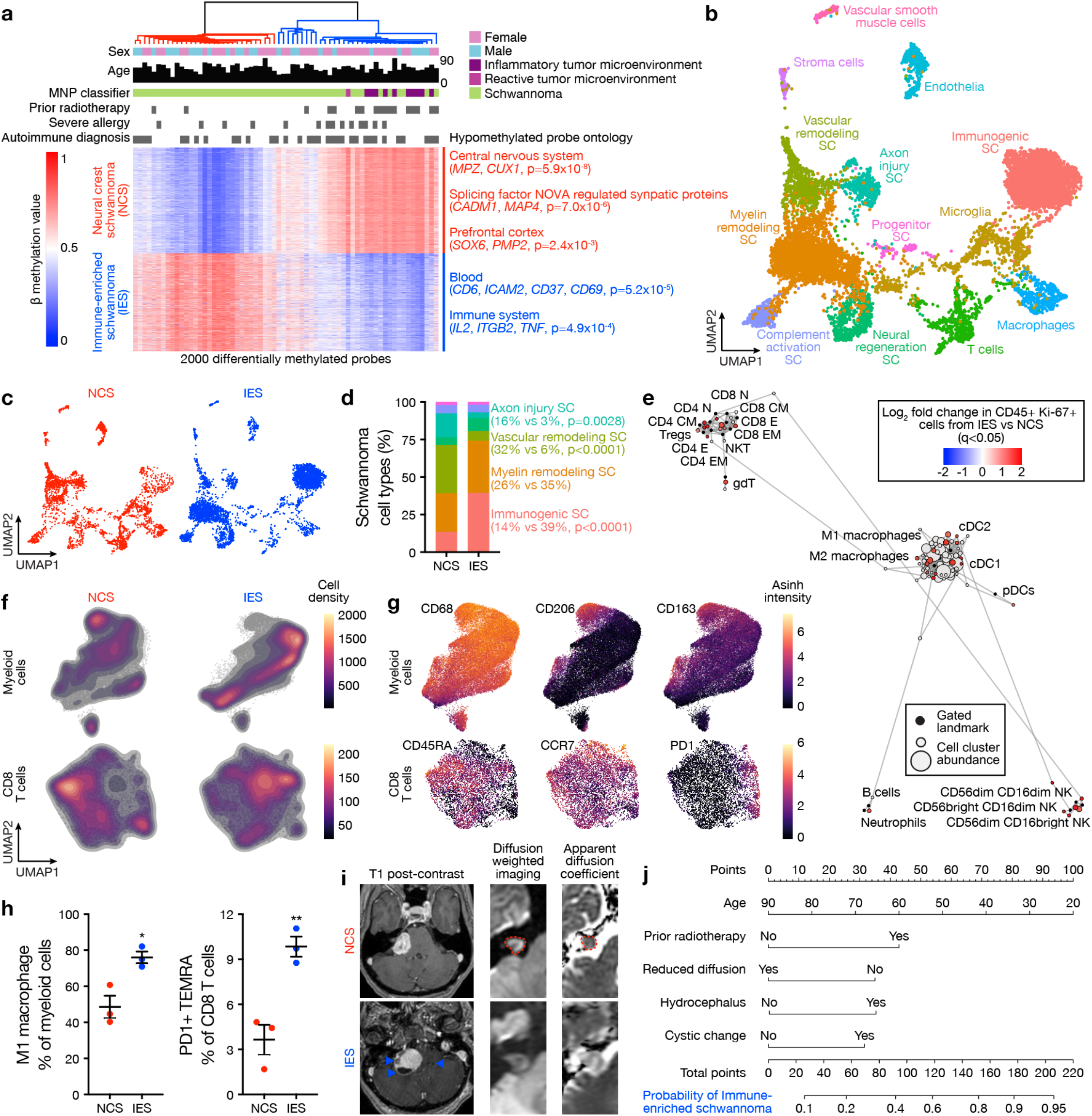
Schwannomas are comprised of neural crest and immune-enriched molecular groups. **a**, Hierarchical clustering of the top 2000 most differentially methylated probes from 66 vestibular schwannomas. Significant gene ontology terms corresponding to hypomethylated probes in neural crest schwannomas (NCS) or immune-enriched schwannomas (IES), clinical metadata, and the molecular neuropathology (MNP) DNA methylation classification of central nervous system tumors^18^ are shown. **b**, Integrated UMAP of 10,628 transcriptomes from harmonized schwannoma single-nuclei (n=6) or single-cell (n=3) RNA sequencing showing schwannoma cell (SC) types and tumor microenvironment cell types. **c**, UMAPs from **b** with individual transcriptomes split according to molecular group of origin. **d**, Relative composition of SC types according to molecular group of origin, colored as in **b. e**, Scaffold plot comprised of 375,355 immune cells from NCS (n=3) or IES (n=3) analyzed using mass cytometry time-of-flight (CyTOF). Manually gated landmark immune cell populations (black) are annotated. Schwannoma immune cell cluster are colored when the proportion of cells are statistically different between IES (positive) and NCS (negative). **f**, CyTOF UMAPs of CD45+ immune cells with overlaid density plots for manually gated myeloid cells (top, 40,000 cells) or CD8 T cells (bottom, 6,632 cells) in NCS (left) or IES (right). **g**, UMAP feature plots of marker genes used to define myeloid or CD8 T cells in **f. h**, CyTOF proportion of schwannoma myeloid cells (left) or CD8 T cells (right) corresponding to M1 macrophages or PD1+ TEMRA CD8 T cells, respectively, in NCS or IES. Lines represent means and error bars represent standard error of means (Student’s t tests, *p:≤0.05, **p:≤0.01). **i**, Representative preoperative magnetic resonance imaging of 66 vestibular schwannomas using T1 post-contrast, T2 diffusion weighted, or apparent diffusion coefficient (ADC) sequences reveals NCS present as solid masses with reduced diffusion (dotted line) and IES present with cystic changes (arrows) and hydrocephalus. **j**, Nomogram for schwannoma immune enrichment and molecular grouping based on non-invasive clinical and magnetic resonance imaging features.

**Fig. 2.**
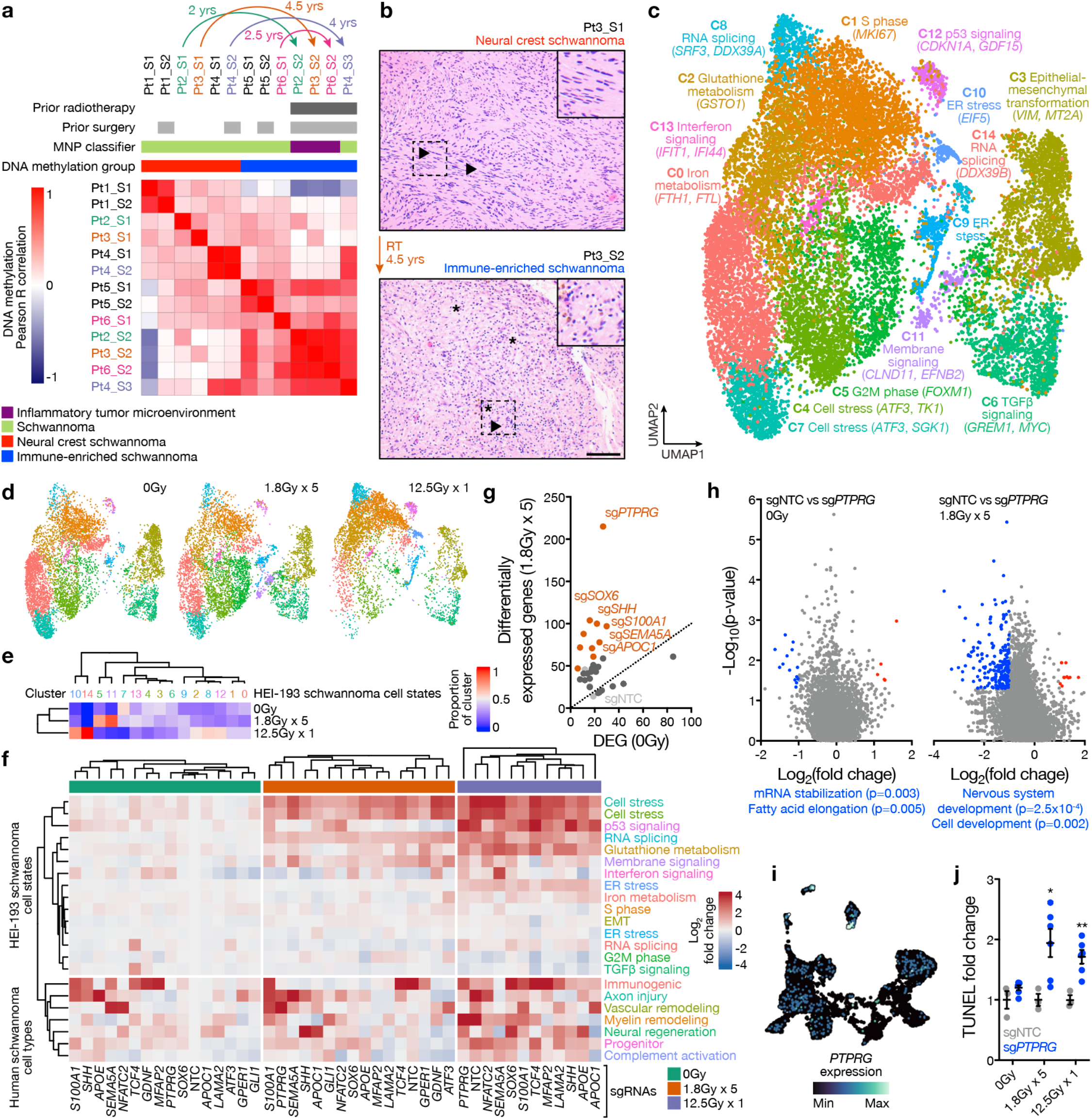
Radiotherapy is sufficient for epigenetic reprogramming of neural crest to immune-enriched schwannoma. **a**, Pairwise Pearson correlation coefficients grouped by hierarchical clustering of DNA methylation profiles from patient (Pt) matched primary and recurrent schwannomas (n=13). Arrows represent reprogramming of primary NCS to IES at recurrence after treatment with radiotherapy. Clinical metadata and the molecular neuropathology DNA methylation classification of central nervous system tumors (MNP)^18^ are shown. **b**, H&E-stained sections of a patient-matched primary NCS and recurrent IES, showing typical histology with Verocay bodies (top arrows) that were replaced by abundant lymphocytes (bottom arrows) with foamy and hemosiderin-filled macrophages (asterisks). RT, radiotherapy. Scale bar, 100μm. **c**, UMAP of 38,754 transcriptomes from single-cell RNA sequencing of HEI-193 cells after treatment with 0Gy (n=3), 1.8Gy x 5 (n=3), or 12.5Gy x 1 (n=3) of radiotherapy revealing 15 distinct schwannoma cell states. **d**, UMAPs from **c** with individual transcriptomes split according to triplicate treatment conditions. **e**, Relative composition of cell states from **c** according to triplicate treatment conditions. **f**, Perturb-seq gene expression heatmap of pseudobulked transcriptomes from HEI-193 cells following sgRNA perturbations (columns) across treatments conditions. Transcriptome modules (rows) are derived from schwannoma cell states in **c** or human schwannoma cell types (Fig. 1b). Expression values are normalized to cells harboring non-targeting control sgRNAs (sgNTC) with 0Gy. **g**, Number of differentially expressed genes (DEG) from schwannoma cell Perturb-seq with radiotherapy (y-axis) versus 0Gy (x-axis). sgRNA perturbations with ≥40 DEG after radiotherapy compared to control are orange. sgRNA not meeting this threshold are dark grey. sgNTCs are light grey. **h**, Volcano plots for differential gene expression analysis from Perturb-seq using MAST for *PTPRG* perturbation compared to sgNTC in 0Gy (left) versus 1.8Gy x 5 conditions (right). Significant positive (red) or negative (blue) gene expression changes are colored (p<0.05, |log_2_(fold change)|>1), corresponding to gene ontology terms. **i**, Feature plot of integrated UMAP from harmonized schwannoma single-nuclei and single-cell RNA sequencing (Fig. 1b) showing *PTPRG* expression in schwannoma cells. **j**, TUNEL staining for apoptosis in HEI-193 cells following CRISPRi suppression of *PTPRG* versus sgNTC after treatment with 0Gy, 1.8Gy x 5, or 12.5Gy x 1 of radiotherapy. Fold changes are normalized to sgNTC in each treatment condition. Lines represent means and error bars represent standard error of means (Student’s t tests, *p:≤0.05, **p:≤0.01).

**Fig. 3.**
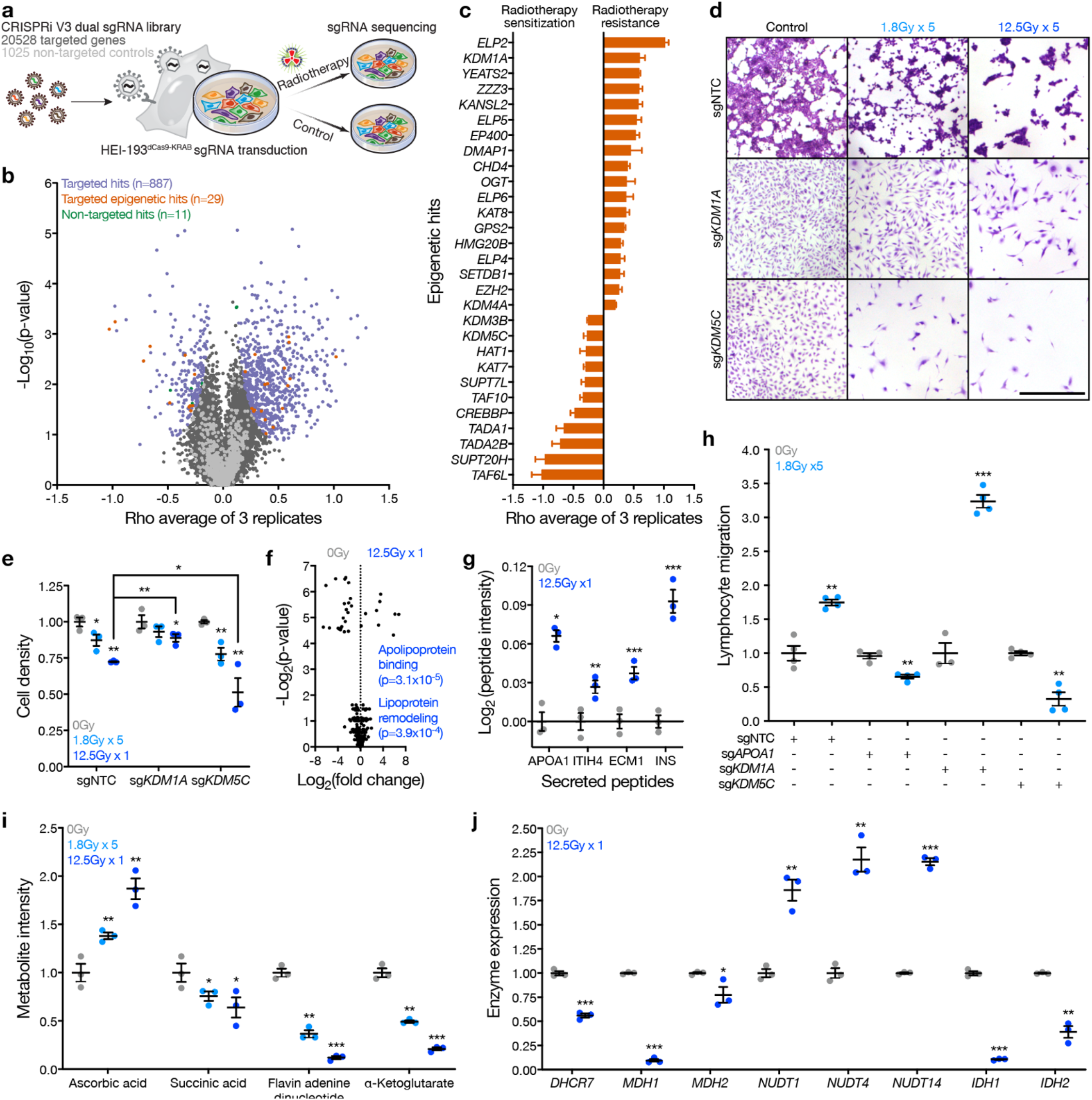
Epigenetic regulators reprogram schwannoma cells and drive immune cell infiltration in response to radiotherapy. **a**, Experimental workflow for triplicate genome-wide CRISPRi screens using dual sgRNA libraries comprised of 20,528 targeted sgRNAs and 1026 non-targeted control sgRNAs (sgNTC). Libraries were transduced into HEI-193 cells that were subsequently treated with 0Gy or 1.8Gy x 5 radiotherapy (n=3 per condition). sgRNA barcodes were sequenced and quantified as proxies for cell enrichment or depletion. **b**, Volcano plot of CRISPRi screen results showing the average rho log_2_(sgRNA in radiotherapy conditions / sgRNA in control conditions) across 3 replicates. On-target hit genes (purple), epigenetic regulator hit genes (orange), and sgNTCs called as hits (green) at a false discovery rate of 1% are shown. **c**, Rho phenotypes of epigenetic regulator CRISPRi screen hits genes from **b. d**, Representative crystal violet staining of HEI-193 cells following CRISPRi suppression of *KDM1A* or *KDM5C* compared to sgNTC after treatment with 0Gy, 1.8Gy x 5, or 12.5Gy x 1 of radiotherapy. Scale bar, 100μm. **e**, Quantification of HEI-193 cell density from **d. f**, Volcano plot of 425 peptides identified using proteomic mass spectrometry of conditioned media from triplicate HEI-193 cultures after radiotherapy or control treatment. Significant gene ontology terms of enriched peptides after radiotherapy conditions are annotated. **g**, Proteomic mass spectrometry parallel reaction monitoring targeted assay validating secreted peptide enrichment in conditioned media from triplicate HEI-193 cultures after radiotherapy as in **f. h**, Transwell primary human peripheral blood lymphocyte migration assays using conditioned media from HEI-193 cells following CRISPRi suppression of *APOA1, KDM1A, or KDM5C ±* radiotherapy as a chemoattractant. **i**, Targeted metabolite mass spectrometry of triplicate HEI-193 cultures after treatment with 0Gy, 1.8Gy x 5, or 12.5Gy x 1 of radiotherapy. Fold changes normalized to 0Gy treatment for each metabolite. **i**, Metabolic enzymes gene expression changes from bulk RNA sequencing of HEI-193 cells *±* radiotherapy (Supplementary Table 6). Fold changes normalized to 0Gy treatment for each gene. Lines represent means and error bars represent standard error of means (Student’s t tests, *p:≤0.05, **p:≤0.01, ***p:≤0.0001).

**Fig. 4.**
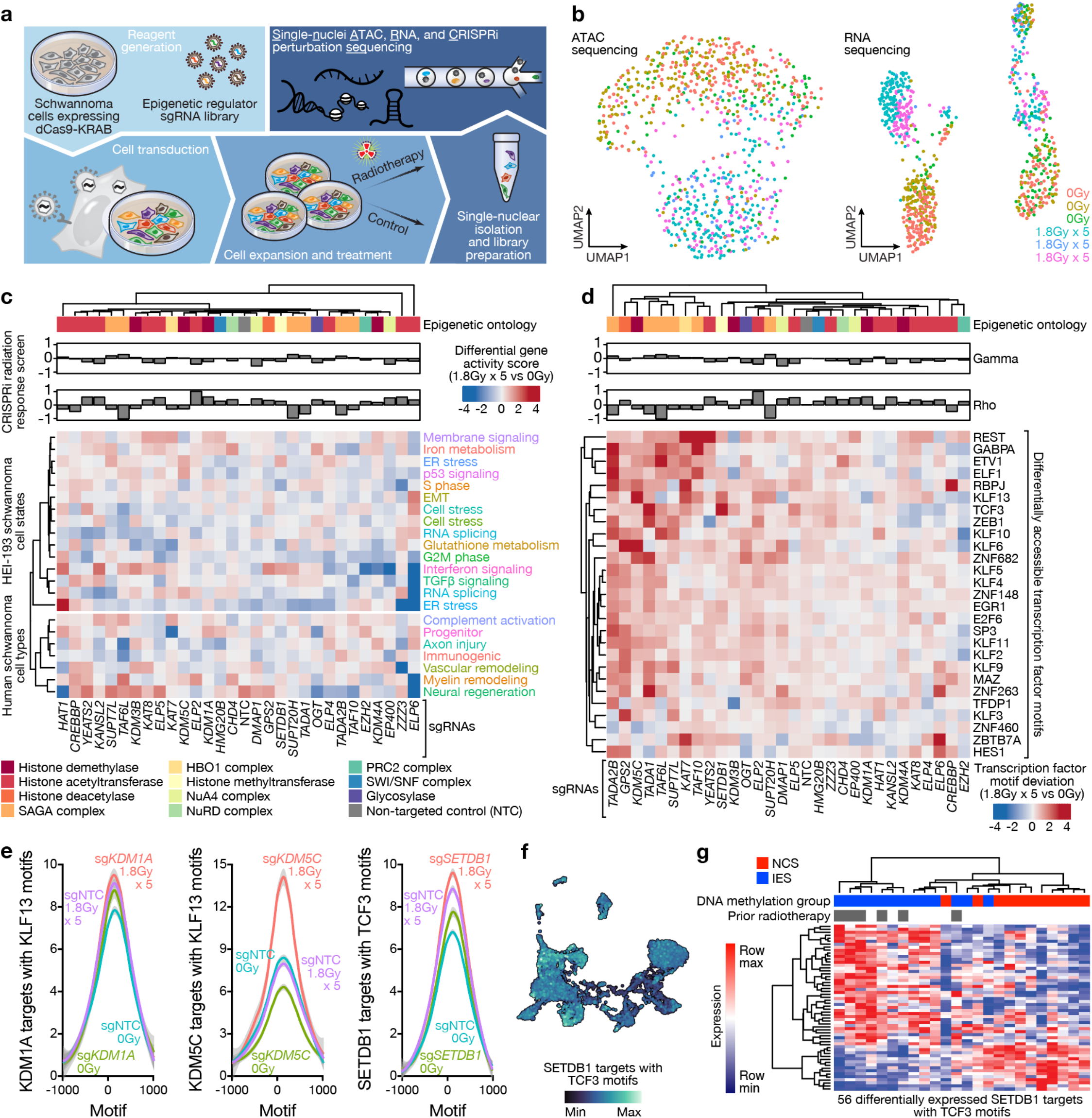
Single-nuclei ATAC, RNA, and CRISPRi perturbation sequencing identifies integrated genomic mechanisms driving schwannoma cell state evolution. **a**, Experimental workflow for <u>s</u>ingle-<u>n</u>uclei <u>A</u>TAC, <u>R</u>NA, and <u>C</u>RISPRi perturbation <u>seq</u>uencing (snARC-seq). Triplicate HEI-193 cultures were transduced with sgRNA libraries targeting 29 epigenetic regulators driving radiotherapy responses from genome-wide CRISPRi screens (Fig. 3c) and treated with 0Gy or 1.8Gy x 5 of radiotherapy prior to isolation of single-nuclei for sequencing. sgRNA identities were recovered from CROP-seq tags in single-nuclei RNA sequencing data. **b**, UMAPs of ATAC (left) or RNA (right) sequencing of 855 single nuclei passing snARC-seq quality control from triplicate control or radiotherapy conditions (Extended Data Fig. 10). **c**, Hierarchical clustering of differential gene activity scores between radiotherapy and control conditions for each snARC-seq perturbation (columns). Gene activity modules (rows) were derived from HEI-193 schwannoma cell states *±* radiotherapy (Fig. 2c) or from human schwannoma cell types (Fig. 1b). Gene ontology of perturbed epigenetic regulators and CRISPRi screen growth (gamma) or radiation response (rho) phenotypes from genome-wide CRISPRi screens (Fig. 3c) are shown. **d**, Hierarchical clustering of differential ChromVAR transcription factor motif deviations between radiotherapy and control conditions for each snARC-seq perturbation (columns). **e**, Average profile plots of normalized ATAC signal at *KLF13* or *TCF3* motifs with ENCODE ChIP-seq peak annotations and differential accessibility following snARC-seq perturbation of *KDM1A, KDM5C*, or *SETDB1*. **f**, Feature plot of integrated UMAP from harmonized schwannoma single-nuclei and single-cell RNA sequencing (Fig. 1b) showing genes near TCF3 motifs that are differentially accessible following *SETDB1* snARC-seq perturbation. **g**, Hierarchical clustering of human schwannoma RNA sequencing profiles using 56 differentially expressed SETDB1 targets with TCF3 motifs showing separation of NCS and IES molecular groups of schwannomas.

**Fig. 5.**
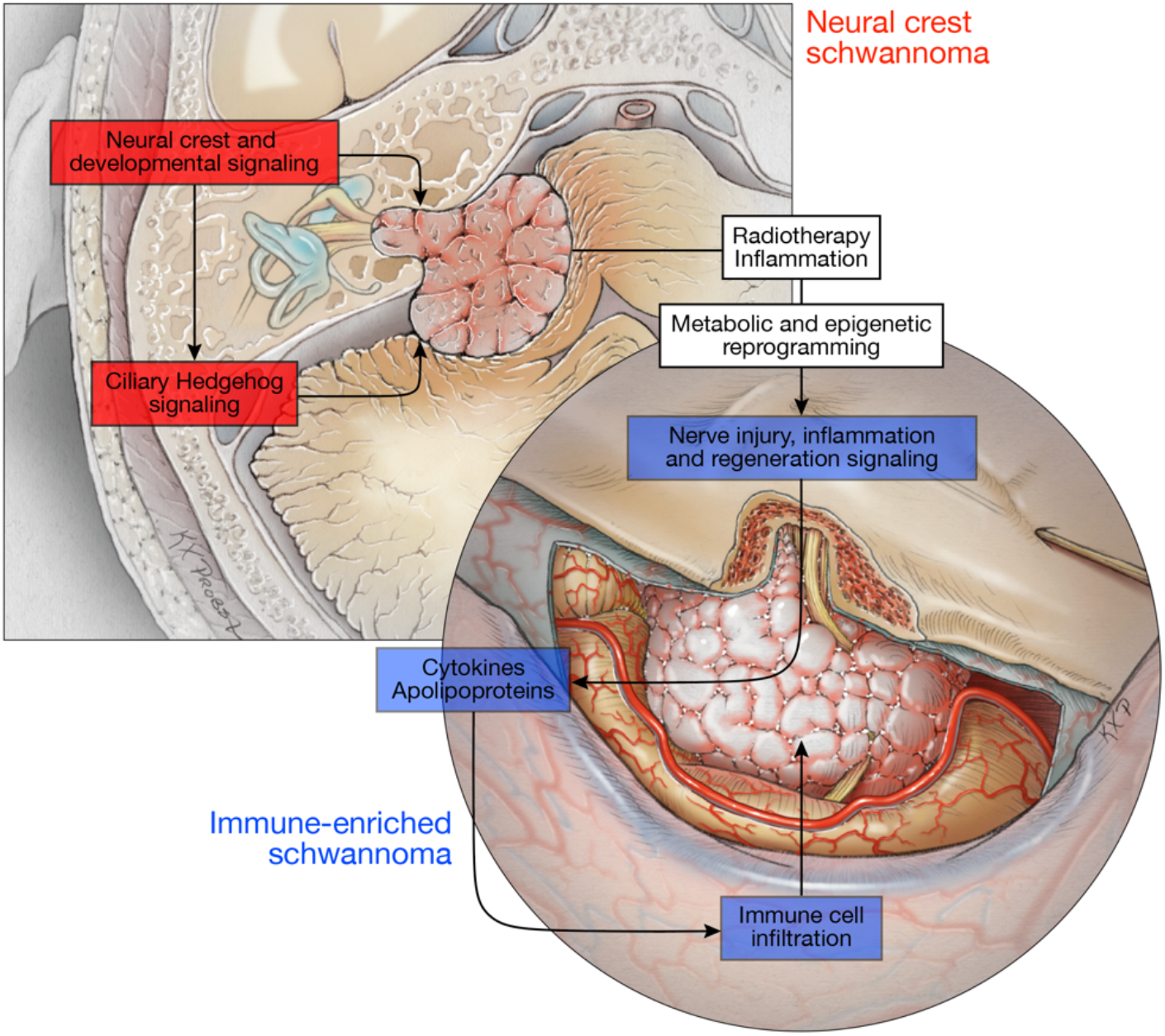
An integrated model of schwannoma tumorigenesis and epigenetic reprogramming.

### Schwannomas are comprised of neural crest and immune-enriched molecular groups

DNA methylation profiling provides robust classification of nervous system tumors and identifies biological drivers of tumorigenesis and response to treatment^18,19^. To define a molecular architecture for mechanistic interrogation of schwannomas, DNA methylation profiling was performed on 66 sporadic vestibular schwannomas from 59 patients who were treated at the University of California San Francisco (Supplementary Table 1). Unsupervised hierarchical clustering and K-means consensus clustering of differentially methylated DNA probes revealed 2 molecular groups (Fig. 1a and Extended Data Fig. 1a). Differentially methylated DNA probes across schwannoma molecular groups were enriched at genes involved in nervous system development or immunologic signaling (Fig. 1a and Supplementary Table 2). Unsupervised hierarchical clustering of 125 sporadic spinal or vestibular schwannomas from an independent institution also identified 2 molecular groups that were distinguished by differential methylation of nervous system development or immune genes (Extended Data Fig. 1b)^14^. Schwannoma clustering using DNA methylation probes overlapping with activate enhancers during neural crest development recapitulated 2 molecular groups of tumors (Extended Data Fig. 2a, b)^20^. RNA sequencing and differential expression analysis on 11 neural crest and 13 immune-enriched schwannomas integrated with DNA methylation profiles revealed enrichment and hypomethylation of either neural crest genes and Hedgehog target genes, or immune genes and apolipoprotein genes, across the 2 molecular groups of schwannomas (Extended Data Fig. 2c, d and Supplementary Table 3). In support of these findings, immunofluorescence on 49 schwannomas showed the Schwann cell differentiation marker SOX10 was enriched in neural crest compared to immune-enriched tumors (Extended Data Fig. 2e), and histologic assessment of 66 schwannomas showed immune-enriched tumors were distinguished by macrophage and lymphocyte infiltration, hyalinized vessels, and coagulative necrosis (Extended Data Fig. 2f).

To define cell types comprising schwannoma molecular groups, single-nuclei or single-cell RNA sequencing was performed on 4 neural crest schwannomas and 5 immune-enriched schwannomas (Extended Data Fig. 3a). Harmonization of 10,628 single-nuclei or single-cell transcriptomes revealed 13 cell clusters in uniform manifold approximation and projection (UMAP) space (Fig. 1b). Expression of the Schwann cell differentiation markers *S100B* and *SOX10* were used to identify 7 clusters of schwannoma cells and 6 clusters of microenvironment cells (Extended Data Fig. 3b). Schwannoma and microenvironment cell types were defined using differentially expressed marker genes (Extended Data Fig. 3c and Supplementary Table 4), unbiased gene expression signatures (Supplementary Fig. 1), and genes associated with active enhancers during demyelinating nerve injury (Extended Data Fig. 3d)^21^. Schwannoma cell types reflected different stages of nerve injury and regeneration (Fig. 1b), which involves axon injury, progenitor cell proliferation, myelin remodeling, vascular remodeling, complement activation, immune recruitment, and neural regeneration^22^. Comparison of cancer cell types in immune-enriched versus neural crest schwannomas showed enrichment of immunogenic schwannoma cells expressing *BCL1* and *CD74*, a regulator of macrophage migration^23,24^, suppression of vascular remodeling schwannoma cells expressing *ADGRB3*, a regulator of angiogenesis and cell proliferation^25–27^; and suppression of axon injury schwannoma cells expressing *NRXN3*, a regulator of synapse function^28^ (Fig. 1c, d). In support of these findings, immunohistochemistry and immunofluorescence revealed schwannoma cell BCL1 expression was enriched in immune-enriched compared to neural crest tumors (Extended Data Fig. 3e, f).

To define immune cell types in the schwannoma microenvironment, mass cytometry by time-of-flight (CyTOF) analysis was performed on 375,355 cells from 3 neural crest schwannomas and 3 immune-enriched schwannomas (Supplementary Table 5). CD45+ immune cells were visualized *in toto* using Scaffold maps^29^ (Fig. 1e and Extended Data Fig. 4a). Proliferating immune cells adjacent to landmark myeloid or lymphoid clusters were enriched in immune-enriched compared to neural crest schwannomas, including memory and effector CD4 and CD8 T cells, γδT cells, NK cells, macrophages, and conventional and plasmacytoid dendritic cells (Fig. 1e). UMAP and PhenoGraph clustering were used to define myeloid and lymphoid immune cell phenotypes^30^ (Fig. 1f, g, h, Extended Data Fig. 4b, and Supplementary Fig. 2-5). In comparison to neural crest schwannomas, myeloid cells in immune-enriched schwannomas were comprised of macrophages with low expression of CD206 and CD163, reflecting an M1 polarization phenotype that is associated with anti-tumor immunity. PD1+ TEMRA CD8 T cells, which are associated with differentiated effector function and tumor control^31,32^, were also enriched in immune-enriched compared to neural crest schwannomas. In support of these phenotypes, immunohistochemistry for the myeloid or lymphoid markers CD68, CD3, and PD1 were enriched in immune-enriched compared to neural crest schwannomas (Extended Data Fig. 4c).

To determine if schwannoma molecular groups could be distinguished using non-invasive biomarkers, we analyzed magnetic resonance (MR) imaging and clinical features for 31 neural crest and 35 immune-enriched schwannomas. Neural crest schwannomas presented as solid masses with reduced MR diffusion compared to immune-enriched tumors (48% versus 15%, p=0.005, Fisher’s exact test) (Fig. 1i). Immune-enriched schwannomas presented with communicating hydrocephalus (38% versus 15%, p=0.04) and cystic changes (53% versus 27%, p=0.03) and had residual tumor adherent to the brainstem after resection (56% versus 21%, p=0.02, Fisher’s exact tests) even after adjusting for tumor size (OR 5.3, 95% CI 1.4-24.1, p=0.02). Univariate and recursive partitioning analyses of MR imaging and clinical features showed prior schwannoma radiotherapy, cystic changes, and the absence of reduced diffusion were specific for immune-enriched schwannomas (Supplementary Fig. 6a). Logistic regression using these features yielded a model with 79% sensitivity and 77% specificity in distinguishing neural crest and immune-enriched schwannomas (AUC 0.79) (Fig. 1j and Supplementary Fig. 6b). Analysis of clinical outcomes showed local freedom from recurrence was equivalent across schwannoma molecular groups (Supplementary Fig. 6c).

### Radiotherapy is sufficient for epigenetic reprogramming of neural crest to immune-enriched schwannoma

Immune-enriched schwannomas were more common than neural crest schwannomas in patients with autoimmune diseases and severe allergic reactions (25.7% vs 0%, p=0.0025), or in patients with prior schwannoma radiotherapy (42.9% versus 6.5%, p=0.0007, Fisher’s exact tests), which occurred a median of 3 years before salvage resection (Fig. 1a and Supplementary Table 1). Analysis of DNA methylation profiles across clinical features revealed >99% of differentially methylated DNA probes in schwannomas could be attributed to prior radiotherapy (Extended Data Fig. 5a). Hierarchical clustering of differentially methylated DNA probes distinguishing schwannomas with prior radiotherapy showed hypomethylation of immune signaling genes and hypermethylation of neuronal progenitor maintenance genes^33^ (Extended Data Fig. 5b). Comparison of DNA methylation profiles between paired primary and recurrent schwannomas from 6 patients (Supplementary Table 1) revealed epigenetic reprogramming of recurrent tumors compared to cognate primary tumors in every instance where radiotherapy was delivered between serial surgeries (Fig. 2a). All recurrent schwannomas with prior radiotherapy from this matched set were immune-enriched and clustered together with pairwise Pearson correlation coefficients that were indistinguishable from randomly-selected unpaired schwannomas with prior radiotherapy (Extended Data Fig. 3c). Moreover, primary neural crest schwannomas showing classic histology were infiltrated by lymphocytes and macrophages that persisted for at least 4 years after radiotherapy, until savage resection of paired samples was performed (Fig. 2b and Supplementary Fig. 7).

RNA sequencing and differential expression analysis showed Hedgehog target genes were enriched in schwannomas without prior radiotherapy (Extended Data Fig. 5d and Supplementary Table 6). Thus, we hypothesized Hedgehog signaling may underlie neural crest schwannomas. Hedgehog signals are transduced through primary cilia^34^. Immunofluorescence revealed schwannoma cell cilia were longer in neural crest compared to immune-enriched schwannomas (Extended Data Fig. 6a), and single-cell and single-nuclei RNA sequencing validated Hedgehog target gene expression in schwannoma cells from human tumors (Extended Data Fig. 6b). Primary cilia were also identified on cultured human Schwann cells, which accumulated Smoothened in cilia and activated the Hedgehog transcriptional program in response to recombinant Sonic Hedgehog (SHH) (Extended Data Fig. 6c-e). Schwann cell proliferation was blocked by the Smoothened antagonist vismodegib or radiotherapy (Extended Data Fig. 6f, g), and radiotherapy also reduced ciliary length, attenuated Smoothened accumulation in primary cilia, and blocked Hedgehog target gene expression (Extended Data Fig. 6h-j). To study these mechanisms in human schwannoma cells we used the HEI-193 cell line, which encodes heterozygous loss of *NF2* on chromosome 22q and a distal splice site mutation in the remaining *NF2* allele^35,36^. CRISPRi suppression of *NF2* in HEI-193 cells stably expressing dCas9-KRAB promoted cell proliferation compared to HEI-193 cells expressing non-targeted control sgRNAs (sgNTC) (Extended Data Fig. 6k, l), suggesting *NF2* retains tumor suppressor functionality in HEI-193 cells. Radiotherapy blocked HEI-193 Hedgehog target gene expression (Extended Data Fig. 6m). In human tumors, the intraflagellar transport gene *IFT88*, which is necessary for assembly of primary cilia, was hypermethylated and suppressed in immune-enriched compared to neural crest schwannomas (Extended Data Fig. 6n, o).

To more broadly define schwannoma cell responses to ionizing radiation, HEI-193 schwannoma cells were treated with fractionated (1.8Gy x 5 fractions) or hypofractionated (12.5Gy x 1 fraction) radiotherapy, both of which are used to treat human schwannomas. Radiotherapy blocked the growth of schwannoma cells (Extended Data Fig. 7a, b), and QPCR assessment of genes distinguishing molecular groups of human schwannomas (Fig. 1) showed radiotherapy induced schwannoma cell expression of inflammatory apolipoproteins^37,38^, immediate early genes that drive cell fate decisions^39^, and *SOX6*, an inhibitor of Schwann cell differentiation^40^ (Extended Data Fig. 7c). To determine if these genes promoted schwannoma cell survival, *APOD* or *SOX6* were suppressed using CRISPRi, which attenuated schwannoma cell proliferation compared to cells expressing sgNTC (Extended Data Fig. 7d, e). DNA methylation profiling and RNA sequencing revealed hypomethylation and enriched expression of immune and apolipoprotein genes in surviving schwannoma cells after radiotherapy (Extended Data Fig. 7f, g). To determine if radiotherapy reprograms schwannoma cell states, single cell RNA-sequencing was performed on HEI-193 cells after control, fractionated radiotherapy, or hypofractionated radiotherapy treatments (Extended Data Fig. 7h). Clustering of 39,569 single-cell transcriptomes revealed 15 cell clusters in UMAP space (Fig. 2c). Schwannoma cell states were defined using differentially expressed marker genes (Extended Data Fig. 7i and Supplementary Table 7), revealing differential activation of diverse cellular mechanisms such as endoplasmic reticulum (ER) stress, interferon signaling, membrane signaling, and p53 signaling (Fig. 2c-e).

To define how schwannoma cell states (Fig. 2c) or cell types (Fig. 1b) are established and respond to radiotherapy, we used Perturb-seq, a functional genomic approach coupling CRISPRi screening with transcriptomic phenotypes in single cells^41–43^. Perturb-seq suppression of 15 genes distinguishing molecular groups of human schwannomas or schwannoma cell states was performed in HEI-193 cells with control, fractionated radiotherapy, or hypofractionated radiotherapy treatments (Supplementary Table 8). Integration of 3,546 single-cell transcriptomes revealed intercellular heterogeneity that was dependent on radiotherapy dose and the cell cycle (Extended Data Fig. 8a-e). We then asked whether Perturb-seq gene suppression affected gene modules distinguishing the 15 schwannoma cell states from single-cell RNA sequencing of schwannoma cells (Fig. 2c), or the 7 schwannoma cell types reflecting different stages of nerve injury and regeneration from integrated single-nuclei and single-cell RNA sequencing of human schwannomas (Fig. 1b). Radiotherapy dose was the primary determinant of cell stress, p53, RNA splicing, and glutathione pathway activation, but genetic perturbations caused heterogenous changes in gene module expression with and without ionizing radiation (Fig. 2f). To identify perturbations disrupting gene expression programs in combination with radiotherapy, differential expression analysis was performed on single cells expressing targeted versus non-targeted sgRNAs with or without radiotherapy (Supplementary Table 9). Suppression of *SHH, SOX6*, or the receptor tyrosine phosphatase *PRPTG*, which regulates neural developmental signaling^44^, exhibited greater numbers of differentially expressed genes after radiotherapy compared to control treatment (Fig. 2g, h and Extended Data Fig. 8f). Integrated single-nuclei and single-cell RNA sequencing showed *PTPRG* was enriched in schwannoma cells from human tumors (Fig. 2i), Perturb-seq suppression of *PTPRG* in schwannoma cells inhibited genes underlying nervous system development only when combined with radiotherapy (Fig. 2h), and CRISPRi suppression of *PTPRG* enhanced schwannoma cell apoptosis with radiotherapy (Fig. 2j and Extended Data Fig. 8g, h). In sum, these data reveal polygenic mechanisms underlie reprograming of neural crest to immune-enriched schwannoma and suggest a role for *PTPRG* in schwannoma response to radiotherapy.

### Epigenetic regulators reprogram schwannoma cells and drive immune cell infiltration in response to radiotherapy

To more broadly define genomic drivers specifying schwannoma cell states in response to ionizing radiation, triplicate genome-wide CRISPRi screens were performed using dual sgRNA libraries comprised of 20,528 targeted sgRNAs and 1026 sgNTCs (Fig. 3a). The effect of genetic perturbations on HEI-193 schwannoma cell growth *±* radiotherapy was defined using targeted DNA sequencing and quantification of integrated sgRNA barcodes over time (Supplementary Table 10). Gene set enrichment analysis of radiotherapy sensitivity screen hits showed expected changes related to the cell cycle and DNA repair, but also revealed significant enrichment of histone acetyltransferase genes (Extended Data Fig. 9a). Examination of screen hits identified 29 epigenetic regulators associated with radiotherapy resistance or sensitivity phenotypes in schwannoma cells (Fig. 3b, c). In support of these findings, CRISPRi suppression of the histone demethylases *KDM1A* or *KDM5C* validated radiotherapy resistance or sensitivity phenotypes, respectively (Fig. 3d, e and Extended Data Fig. 9b).

Cancer cell death from ionizing radiation leads to acute and often transient recruitment of immune cells to the tumor microenvironment^45^, but we found surviving schwannoma cells existing alongside infiltrating immune cells for many years after treatment of human tumors with radiotherapy (Fig. 2b, Extended Data Fig. 2f, 4c, and Supplementary Fig. 7). To determine if schwannoma cell reprogramming contributes to immune cell infiltration of the tumor microenvironment, proteomic mass spectrometry was performed on conditioned media from surviving HEI-193 schwannoma cells after radiotherapy (Supplementary Fig. 8a-c). Ontology analyses of 425 differentially expressed proteins from conditioned media revealed enrichment of apolipoproteins after ionizing radiation (Fig. 3f), and a parallel reaction monitoring targeted assay validated secretion of APOA1 and other chemokines from surviving schwannoma cells in response radiotherapy (Fig. 3g). Transwell migration assays showed conditioned media from schwannoma cells recruited primary human peripheral blood lymphocytes after radiotherapy (Extended Data Fig. 9c), and conditioned media from schwannoma cells with CRISPRi suppression of the radiotherapy resistance hit *KDM1A* enhanced lymphocyte migration (Fig. 3h). In contrast, conditioned media from schwannoma cells with CRISPRi suppression of *APOA1* or the radiotherapy sensitivity hit *KDM5C* inhibited lymphocyte migration (Fig. 3h and Extended Data Fig. 9b). Thus, epigenetic mechanisms in schwannoma cells contribute to immune cell infiltration in response to radiotherapy.

To determine if altered expression of epigenetic regulators mediates schwannoma responses to radiotherapy, we analyzed the 29 epigenetic hits from our genome-wide CRISPRi screen (Fig. 3b, c) across schwannoma cell types (Fig. 1b) and schwannoma cell states (Fig. 2c). Surprisingly, no epigenetic regulators were differentially expressed across either context, either with or without radiotherapy (Supplementary Table 4,7). Epigenetic regulator activity is dependent on metabolite cofactors that covalently modify histone subunits^46^. Thus, we hypothesized schwannoma cell metabolites may be altered in response to radiotherapy. To test this, we performed targeted metabolite profiling of HEI-193 cells after control, fractionated radiotherapy, or hypofractionated radiotherapy treatments using metabolomic liquid chromatography mass spectrometry. Radiotherapy suppressed the KDM5C cofactor α-ketoglutarate and succinic acid, a biproduct of α-ketoglutarate metabolism (Fig. 3i). α-ketoglutarate was also suppressed by radiotherapy in primary schwannoma cells from 7 patients (Supplementary Fig. 8d). Moreover, radiotherapy suppressed the KDM1A cofactor flavin adenine dinucleotide (FAD) and increased the KDM5C cofactor ascorbic acid (Fig. 3i). RNA sequencing showed *IDH1/2* and *MDH1/2*, which produce α-ketogluterate or succinic acid, respectively, were suppressed following schwannoma cell radiotherapy (Fig. 3j). *DHCR7*, which metabolizes ascorbic acid precursors, was also suppressed after radiotherapy, but the nudix hydrolases *NUTD1/4/14*, which degrade FAD, were increased (Fig. 3j). These results suggest altered expression of metabolic enzymes and metabolite cofactors indirectly influence epigenetic regulator activity during schwannoma radiotherapy.

### Single-nuclei ATAC, RNA, and CRISPRi perturbation sequencing identifies integrated genomic mechanisms driving schwannoma cell state evolution

To define how epigenetic regulators shape chromatin accessibility and gene expression in schwannoma cells during tumor evolution, we developed a technique for simultaneous interrogation of chromatin accessibility and gene expression coupled with genetic and therapeutic perturbations in single-nuclei (Fig. 4a). <u>S</u>ingle-<u>n</u>uclei <u>A</u>TAC, <u>R</u>NA, and <u>C</u>RISPRi perturbation sequencing (snARC-seq) of the 29 epigenetic regulator hits from our genome-wide CRISPRi screen (Fig. 3b, c) was performed in HEI-193 cells with control or radiotherapy treatments (Extended Data Fig. 10a). sgRNA identities were assigned to individual cells using a hypergeometric test (Extended Data Fig. 10b, c). Genome-wide ATAC signals were enriched at transcription start sites and 5’ nucleosome free regions^47^ (Extended Data Fig. 10d). ATAC and RNA sequencing data exhibited heterogenous distributions in UMAP space (Fig. 4b). Together, these data support successful simultaneous profiling of 3 genomic modalities in single-nuclei with or without radiotherapy (Fig. 4a).

To determine if snARC-seq suppression of chromatin regulators reprogrammed the epigenetic landscape of schwannoma cells, we quantified gene activity scores for open chromatin regions nearby marker genes distinguishing schwannoma cell states (Fig. 2c) or cell types (Fig. 1b). Differential gene activity scores clustered according to gene ontologies of perturbed epigenetic regulators (e.g. histone demethylases, histone acetyltransferases) and CRISPRi screen phenotypes (radiotherapy sensitivity, negative rho; radiotherapy resistance, positive rho) (Fig. 4c). For instance, snARC-seq suppression of elongator component *ELP6* attenuated interferon signaling and ER stress schwannoma cell states that were normally activated by radiotherapy (Fig. 2c-e), consistent with the role of the elongator acetyltransferase complex in gene activation and sensitivity to genotoxic stress^48,49^.

To elucidate transcription factors underlying changes in gene expression programs with snARC-seq suppression of chromatin regulators, we scored transcription factor motif deviation with or without radiotherapy^50^ (Fig. 4d). Restricting analysis to transcription factors expressed in HEI-193 cells (Supplementary Table 6) revealed heterogenous disruptions in chromatin accessibility that clustered based on CRISPRi screen phenotypes (Fig. 4d). For instance, snARC-seq suppression of the radiotherapy sensitivity hit *KDM5C* caused enrichment of open chromatin regions at KLF13 motifs with radiotherapy (Fig. 4e), and KLF13 target genes such as *DDI2* and *PTPA* were simultaneously accessible at their genomic loci and had enriched mRNA expression in single-nuclei following snARC-seq suppression of *KDM5C* with radiotherapy (Extended Data Fig. 10e). *ITIH4*, a secreted protein that was enriched in conditioned media from schwannoma cells surviving radiotherapy (Fig. 3g), displayed closed chromatin and RNA suppression following snARC-seq suppression of *KDM5C* with radiotherapy (Extended Data Fig. 10e). Concordant changes in KLF13 gene activity were not observed following snARC-seq suppression of the radiotherapy resistance hit *KDM1A* with radiotherapy (Fig. 4e), consistent with opposing phenotypes of *KDM1A* and *KDM5C* in mediating schwannoma radiotherapy responses (Fig. 3c). snARC-seq suppression of the histone methyltransferase *SETDB1* increased accessibility at TCF3 motifs and expression of TCF3 target genes such as *RNF217, FOXO1*, and *RAD23B* with radiotherapy (Extended Data Fig. 10e). Integrated single-nuclei and single-cell RNA sequencing showed TCF3 target genes were expressed in schwannoma cells from human tumors (Fig. 4f), and schwannoma clustering restricted to RNA sequencing of TCF3 target genes recapitulated neural crest and immune-enriched molecular groups of schwannomas (Fig. 4g). Thus, our snARC-seq technique for simultaneous interrogation of chromatin accessibility and gene expression coupled with genetic and therapeutic perturbations in single-nuclei reveals epigenetic dependances underlying radiotherapy responses in schwannomas cells that are conserved across molecular groups of human tumors.

## Discussion

Mutational reprogramming and non-mutational epigenetic reprogramming are critical for cancer evolution^1,12^. Schwannomas are the most common tumors of the peripheral nervous system and have a low burden of somatic mutations that is not increased by radiotherapy^5,14,15^. Our results reveal schwannomas are comprised of neural crest and immune-enriched molecular groups, and that radiotherapy is sufficient for epigenetic reprogramming of neural crest to immune-enriched schwannoma (Fig. 5). Thus, schwannomas represent prototypical cancers for understanding epigenetic mechanisms specifying cancer cell states and intratumor heterogeneity. To that end, our results also establish a framework for defining how epigenetic reprogramming modulates cancer evolution in response to treatment. We integrate multiplatform profiling of human tumors with multiomic functional genomic approaches to show interconversion of schwannoma molecular groups in response to an oncologic treatment that is used for half of cancer patients worldwide^51^. Moreover, we develop a new method for simultaneous profiling of epigenetic and transcriptional cell states following genetic and therapeutic perturbations in single-nuclei (snARC-seq), a technique that may more broadly enable fundamental discoveries in health and disease.

## Acknowledgements

The authors thank Jeremy Reiter, Monika Sigg, Aparna Bhaduri, and Aaron Tward for providing comments and reagents; Eric Chow, Derek Bogdanoff, Tomasz Nowakowski, Chang Kim, and Galina Schmunk for technical sequencing assistance and analysis; and Ken Probst and Noel Sirivansanti for Illustrations. This study was supported by the ASCO Conquer Cancer Sontag Foundation Young Investigator Award to SJL; NIH grants R21 HD106238 and R01 CA251221, and DOD grant NFRP NF200021, to DRR; NIH grant U54 CA209891 to NJK; and a Children’s Tumor Foundation Young Investigator Award to HNV.

## Author contributions statement

All authors made substantial contributions to the conception or design of the study; the acquisition, analysis, or interpretation of data; or drafting or revising the manuscript. All authors approved the manuscript. All authors agree to be personally accountable for individual contributions and to ensure that questions related to the accuracy or integrity of any part of the work are appropriately investigated, resolved, and the resolution documented in the literature. SJL, NWC, MP, MCD, JEVM, DLS, HNV, UEL, CDE, NJK, MH, MHS, LG, PVT, and DRR designed the study and analyses. Experiments were performed by SJL, TCC, NWC, JS, MP, MCD, KF, WCC, DLS, HNV, AC, JDB, UEL, KJHG, ES, KHC, BVL, DW, and DRR. Data analysis was performed by SJL, TCC, NWC, JS, MP, MCD, WCC, JEVM, DLS, HNV, UEL, and DRR. The study was supervised by SJL, HNV, SEB, PKS, STM, DL, MWM, MSB, AP, NJK, MH, MHS, LG, PVT, and DRR. The manuscript was prepared by SJL and DRR with input from all authors.

## Competing interests statement

The authors declare no competing interests.

## Tables

Not applicable

## Methods

### Vestibular schwannomas

Patients presenting for resection of sporadic vestibular schwannoma who gave consent for tumor sampling for research were included in the retrospective discovery cohort or in single-cell RNA sequencing and CyTOF experiments (Supplementary Table 1). This study was approved by the UCSF Institutional Review Board (#10-01318, #18-24633). Exclusion criteria included a history of neurofibromatosis type 2 and, with the exception of prospectively-obtained schwannomas for single-cell RNA sequencing and CyTOF experiments, less than 18 months of magnetic resonance imaging follow-up. Vestibular schwannomas for DNA methylation profiling, bulk RNA sequencing, and single-nucleus RNA sequencing were identified from the UCSF Brain Tumor Center Biorepository & Histology Core, and all samples from patients meeting inclusion and exclusion criteria were included, resulting in 76 clinically heterogeneous retrospective and prospective schwannomas from 69 patients who were treated from 2003 to 2020. Of note, many patients in this cohort were initially treated with hybrid strategies employing primary tumor debulking followed by adjuvant radiotherapy in an attempt to optimize outcomes^52,53^. However, such strategies are now known to be associated with higher recurrence rates and are not recommended. All cases with available material were reviewed by a board-certified neuropathologist for diagnostic confirmation (MP). Additional clinical variables were analyzed by chart review (SJL, JDB, DRR).

### Annotation of radiologic features

A comprehensive radiologic review of preoperative magnetic resonance imaging studies for the patients comprising the retrospective discovery cohort (Supplementary Table 1) was performed by a board-certified neuroradiologist who was blinded to all clinical and molecular data (JEVM). Anatomic magnetic resonance images including T2, T2 fluid attenuated inversion recovery (FLAIR), and post-contrast T1 weighted images were reviewed. Diffusion weighted imaging (DWI) including derived apparent diffusion coefficient (ADC) maps were also evaluated. Radiologic features of enhancement pattern (homogeneous or heterogeneous), cystic change, mass effect, peritumoral edema, indistinct brain-tumor interface, lobulated margins, and hydrocephalus were scored in categoric binary fashion and were not found to be significant different between molecular groups of schwannomas unless stated otherwise. Hydrocephalus was further characterized as non-communicating or communicating. Diffusion characteristics were assessed qualitatively (reduced or facilitated) in vestibular schwannomas for which diffusion weighted imaging was acquired.

### DNA methylation profiling and analysis

Genomic DNA was isolated from schwannomas by mechanical homogenization (Qiagen TissueLyzer) followed by the All-Prep Universal Kit for DNA and RNA isolation (Qiagen). DNA quality was tested by spectrophotometry and clean-up was performed using DNA precipitation as needed. Genome-wide methylation profiles were obtained with the Illumina MethylationEPIC 850K array for each schwannoma. Data preprocessing and normalization was performed using *minfi* version 1.30 in R version 3.6.0^54,55^. Methylation probes with detection significance p>0.05 were excluded from analysis. Normalization was performed using functional normalization^56^. Probes were filtered according to the following criteria: (i) exclusion of probes mapping to sex chromosomes (n=11,551), (ii) exclusion of probes containing a common single nucleotide polymorphism (SNP) within the targeted CpG site or on an adjacent base pair (n=24,536), and (iii) exclusion of probes not mapping uniquely to the human reference genome hg19 (n=9,993). A total of 815,630 probes were retained for further analysis. Methylation β values were calculated as the ratio of methylated probe intensity to the sum of methylated and unmethylated probe intensities and visualized on heatmaps in an absolute scale^57^. Variable probes were identified by ranking all methylation probes by β value variance across all schwannomas. The top 2000 probes by variance were then subjected to consensus clustering using complete linkage hierarchical clustering with 1-Pearson correlation as distance matrix, sampling between 2-6 possible clusters with 20 resampling iterations. Schwannomas belonging to each of the 2 molecular groups by hierarchical clustering were used for differential methylation analysis against methylation β values using the dmpFinder function in *minfi*. The resulting differentially methylated probes with FDR<0.05 were ranked by log odds values between the 2 molecular groups, and the top and bottom 1000 differentially methylated probes (total 2000) were used for hierarchical clustering of schwannomas. Gene ontology analysis of probes was performed by extracting the associated gene body annotation from each probe using the Bioconductor package IlluminaHumanMethylationEPICanno.ilm10b2.hg19 and inputting the gene lists grouped based on the results of hierarchical clustering into Enrichr^58^.

To compare DNA methylation with RNA sequencing expression data, we used all methylation probes corresponding to each gene as annotated in the Illumina MethylationEPIC manifest, which includes transcription start site (TSS), gene body, and 3’/5’ UTR probes, in order to interrogate beyond promoter methylation. We ranked the differentially methylated probes from most to least significant according to the FDR and compared the most significant probe for each gene to the expression fold change for each differentially expressed gene in RNA sequencing analysis.

Random forest classification of global methylation profiles was performed to compare each methylation profile with a broader set of CNS malignancies as previously described^18^, revealing that immune enriched schwannomas classified as either inflammatory tumor microenvironment or reactive tumor microenvironment (Fig. 1a), consistent with the significant immune cell infiltration in these tumors. External validation methylation data was obtained from GSE79009^14^ and preprocessed and normalized as above. The overlapping set of DNA methylation probes between the external validation cohort (Illumina Methylation 450K array) and the 2000 differentially methylated probes from the discovery cohort (Illumina MethylationEPIC 850K array) was used for complete linkage hierarchical clustering of external validation cohort schwannomas with 1-Pearson correlation. Differential methylation analysis of clinical variables was performed by separating schwannoma methylation profiles based on binary variables (sex, prior surgery, prior radiotherapy) or bisecting the cohort at the median of continuous variables (size, growth rate, age) and performing differential methylation analysis as described above. DNA methylation probes at neural crest enhancers were identified by overlapping the coordinates of each methylation site with those of active neural crest enhancers in development^59^.

To classify prospective vestibular schwannomas analyzed using single-cell RNA sequencing and CyTOF into molecular groups, a methylation-based classifier using differentially methylated probes from our discovery cohort of 66 tumors was used to construct a support vector machine (SVM) using *caret* version 6.0 in R version 3.6.0. A linear kernel SVM was constructed using training data comprising 75% of randomly selected samples from the discovery cohort with 10-fold cross validation. The top 2000 differentially methylated probes were used as variables. The model was applied to a test set comprising 25% of randomly selected samples from the discovery cohort, and receiver operating characteristics were acquired for 1000x resampling of test data. The SVM distinguished neural crest from immune-enriched schwannomas with 100% accuracy when used to classify randomly selected test sets of samples from our initial cohort (95% CI 78.2-100%, p=8.04×10^−5^).

Copy number variant (CNV) calling was performed from schwannoma DNA methylation profiles, and revealed that loss of chromosome 22q, which contains the tumor suppressor *NF2*, was the most common CNV and occurred in 32% of tumors (Supplementary Table 1). To evaluate whether we could identify single nuclei CNVs from prospective schwannomas that demonstrated loss of chromosome 22q based on DNA methylation profiling, we applied CONICS^60^, but were unable to identify a distinct tumor population harboring 22q loss, likely due, in part, to technical factors related to single-nuclei sequencing data from frozen specimens and minimal chromosomal instability of schwannomas^14,15^. Methylation arrays were deconvolved to identify tumor cell purity and constituent cell types (Supplementary Table 1) based on published approaches enabling comparison to a reference atlas of pure methylation profiles^61^ or with a random forest classifier trained on the ABSOLUTE method developed by the TCGA for whole exome sequencing^62^.

### Fluorescence microscopy

Fluorescence microscopy was performed on a Zeiss LSM 800 confocal laser scanning microscope with Airyscan. Images were processed and quantified from at least 2 regions per condition using ImageJ^63^. Cilia prevalence was quantified as the ratio of cilia to nuclei, and ciliary fluorescence intensity was quantified from regions of interest normalization to background fluorescence.

### Histology and light microscopy

Formalin-fixed and paraffin-embedded whole slide sections were reviewed for histologic features including presence of capsule, cystic degeneration, Verocay bodies, biphasic morphology (Antoni A and Antoni B zones), degenerative atypia, hypercellular zones (bone-to-back or overlapping nuclei), hyalinized/thick-walled intratumor vessels, necrosis, hemosiderin deposition, calcification, hyalinization and mitoses by a board-certified neuropathologist who was blinded to all clinical and molecular data (MP). Mitotic activity was noted in regions of maximal activity as the number of mitotic figures per 10 consecutive high-power fields measuring 0.24 mm^64^. Inflammatory infiltrate including lymphocytes and macrophages were scored on H&E-stained sections as no immune cells, scattered immune cells, or abundant immune cells. All vestibular schwannomas with sufficient tissue were included in tissue microarrays (TMAs) containing 4 μm thick paraffin sections from 2 mm cores in duplicate for each case. In cases with biphasic histology, the cores predominantly targeted the cellular Antoni A zones and avoided areas with near-complete macrophage infiltrate or necrosis. All histologic features and stains were scored for each TMA core separately and were averaged for each case.

### Immunofluorescence and immunohistochemistry

Immunofluorescence for schwannoma SOX10 and cilia expression was performed using whole slide sections and TMAs. Sections were deparaffinized in xylene, rehydrated through graded ethanol dilutions and subjected to antigen retrieval using CC1 TRIS buffer (Ventana Medical Systems); labeled with primary antibodies including SOX10 to mark schwannoma cells (API 3099, Biocare; labeling validated in schwannoma and melanoma), Pericentrin (PA5-54109, Thermo Fisher Scientific; labeling validated by Pericentrin knockdown) and γTubulin (T5192, Sigma; labeling validated in human chicken cells) to mark centrosomes, and Acetylated Tubulin to mark cilia (T6793, Sigma; labeling validated in vertebrate and invertebrate organisms); labeled with Alexa Fluor secondary antibodies and DAPI to mark DNA (62248, Thermo Fisher Scientific); and mounted in mounted in ProLong Diamond Antifade Mountant (Thermo Fisher Scientific).

Immunofluorescence for HSC and MSC cilia was performed on glass coverslips. Cells were fixed in 4% paraformaldehyde, blocked in 2.5% FBS, 200 mM glycine and 0.1% Triton X-100 in PBS for 30minutes at room temperature (Thermo Fisher Scientific), and labeled with Smoothened (ab72130, Abcam; labeling validated by competition with immunizing peptide) and Centriolin (sc-365521, Santa Cruz Biotechnology; labeling validated using THP-1, SK-BR-3 and U-937 cell lysates) primary antibodies at 4°C overnight. Cells were labeled with Alexa Fluor secondary antibodies, Hoescht to mark DNA (H3570, Life Technologies), and Acetylated Tubulin Alexa Fluor 647 Conjugate to mark cilia (sc-23950, Santa Cruz Biotechnology; labeling validated using A2058, 3T3-L1 and Jurkat cell lysates), for 1 hour at room temperature. Cells were mounted in ProLong Diamond Antifade Mountant (Thermo Fisher Scientific).

Immunohistochemical stains for T cells and macrophages were performed using TMAs using standard techniques. In brief, slides were subjected to antigen retrieval and labeled with CD3 (A0452, Agilent Technologies, Santa Clara, CA; labeling validated by over-expression) or CD68 (M0814, Agilent Technologies; labeling validated using human B-cell lymphoma) primary antibodies. Detection was performed using the UltraView Universal DAB kit on the Ventana Benchmark Platform (Ventana Medical Systems). CD3 was scored as the number of positive cells in each core as negative (*≤*10), low (11-50), moderate (51-100) and abundant (>100). CD68 was scored as negative (no positive cells recognizable at high power), weak staining in <10% of cells), moderate (staining in 10-20% of cells) or abundant (staining in >20% of cells, easily visible at low power). Immunohistochemistry for BCL1 (RM9104R7, Thermo Fisher Scientific labeling validated using MAD109 cell lysate) was performed as described for T cells and macrophages but on whole slide sections and with scoring as was done for CD68.

### Cell culture and treatments

Human Schwann cells (HSC) were cultured in complete Schwann Cell Medium on Poly-L-Lysine coated substrates (ScienCell Research Laboratories). HEI-193 schwannoma cells were cultured in Dulbecco’s Modified Eagle Medium supplemented with 10% fetal bovine serum (FBS), glutamine (Thermo Fisher Scientific)^35^. Subconfluent HSC and HEI-193 schwannoma cells were irradiated with a X-Rad 320 (Precision X-Ray) irradiator using a 320 KV output at a rate of 3 Gy/min, with rotating a platform supporting cell culture plates, and cells were quantified by manual hemocytometer during routine cell culture passaging. Crystal violet staining of HEI-193 cells was quantified with background subtraction using ImageJ^63^. For ciliation and Hedgehog signaling assays, cultures were transitioned to OptiMEM (Thermo Fisher Scientific) and treated with recombinant Sonic Hedgehog 1 μg/ml (1845, R&D Systems, Minneapolis, MN) or vehicle control for 24 hours. Subconfluent HSC cultures in 96 well plates were treated with vismodegib (Genentech) for 72 hours and cell proliferation was assayed using the CellTiter 96 Non-Radioactive Cell Proliferation kit and a GloMax Discovery plate reader (Promega). For proteomic mass spectrometry, cells were grown in serum free N5 media consisting of NeuralBasal A supplemented with N2 and B27 supplements without vitamin A, L-glutamine 2 mM, antibiotic-antimycotic (Thermo Fisher Scientific), bFGF 20 ng/mL (119-126, VWR), and human EGF 20 ng/mL (AF-100-15, PeproTech). Cells were irradiated, and cell-free conditioned media was isolated by centrifugation for 10 minutes at 4000g and filtering through a 0.45 μm syringe for mass spectrometry. TUNEL assays for apoptosis were performed using the APO-BrdU™ TUNEL Assay Kit, with Alexa Fluor™ 488 Anti-BrdU (Thermo Scientific).

### CRISPR interference

HEI-193 schwannoma cells stably expressing the CRISPR interference (CRISPRi) components dCas9-KRAB were generated as previously described^65,66^. Briefly, HEI-193 schwannoma cells were transduced with lentivirus harboring SFFV-dCas9-BFP-KRAB, and the top ∼25% of cells expressing BFP were FACS sorted and expanded. For single gene targeted knockdowns, sgRNA protospacer sequences (Supplementary Table 11) were cloned into a lentiviral expression vector (U6-sgRNA EF1Alpha-puro-T2A-BFP) by annealing and ligation. HEI-193 schwannoma CRISPRi cells were then transduced with sgRNA lentivirus and selected with puromycin 1μg/mL for at least 4 days before experimentation.

### Lymphocyte isolation and migration

Peripheral blood lymphocytes were isolated from human blood of healthy volunteers^67^. A Polymorph density gradient (Accurate Chemical & Scientific Corporation) was used to isolate peripheral blood mononuclear cells that were subsequently selected and differentiated into T cell lymphocytes in Roswell Park Memorial Institute 1640 media (Life Technologies) supplemented with 10% FBS, 1% penicillin/streptomycin, phytohemagglutinin 1 μg/ml (10576015, Thermo Fisher Scientific) and recombinant human IL-2 20 ng/ml (202-IL, R&D Systems). The QCM Leukocyte Migration Assay (MilliporeSigma) and a GloMax Discovery plate reader (Promega) were used to quantify transwell T cell lymphocyte migration from the apical chamber over 4 hours at 37°C. Unconditioned HEI-193 media as a negative control, unconditioned HEI-193 media supplemented with recombinant human CCL21 600 ng/ml (366-6C, R&D Systems) as a positive control, HEI-193 media 5 days after 12.5Gy in 1 fraction, or HEI-193 media without radiation were placed in the basolateral chamber. All conditions were free from serum or other supplements unless specifically indicated.

### Mass cytometry by time-of-flight cell preparation and analysis

All mass cytometry by time-of-flight (CyTOF) antibodies and concentrations used for analysis can be found in Supplementary Table 5. Primary conjugates of antibodies were prepared using the MaxPAR antibody conjugation kit (Fluidigm). Following labeling, antibodies were diluted in Candor PBS Antibody Stabilization solution (Candor Bioscience) supplemented with 0.02% NaN3 to between 0.1 and 0.3 mg/mL and stored long-term at 4°C. Each antibody clone and lot were titrated to optimal staining concentrations using primary human samples.

All tissue preparations were performed simultaneously from each sample as previously described^29^. Tumors were finely minced and digested in L-15 medium with 800 units/ml collagenase IV (Worthington) and 0.1 mg/ml DNase I (Sigma). After digestion, re-suspended cells were quenched with PBS/EDTA at 4°C. All tissues were washed with PBS/EDTA and re-suspended 1:1 with PBS/EDTA and 100 mM Cisplatin (Enzo Life Sciences) for 60 seconds before quenching 1:1 with PBS/EDTA/BSA to determine viability^29^. Cells were centrifuged at 500 g for 5 min at 4°C and re-suspended in PBS/EDTA/BSA at a density between 1-10 × 10^6^ cells/ml. Suspensions were fixed for 10 min at RT using 1.6% paraformaldehyde and frozen at -80°C.

Mass-tag cell barcoding was performed as previously described^68^. Briefly, 10^6^ cells from each sample were barcoded with distinct combinations of stable Pd isotopes in 0.02% saponin in PBS. Samples were then barcoded together. Cells were washed once with cell staining media (PBS with 0.5% BSA and 0.02% NaN3), once with 1X PBS, and pooled into a single FACS tube (BD Biosciences). After data collection, each condition was deconvoluted using a single-cell debarcoding algorithm^68^.

Cells were resuspended in cell staining media comprised of PBS with 0.5% BSA and 0.02% NaN3, and antibodies against CD16/32 were added at 20 mg/ml for 5 min at RT on a shaker to block Fc receptors. Surface marker antibodies were then added, yielding 500 uL final reaction volumes and stained for 30 min at room temperature on a shaker. Following staining, cells were washed 2 times with cell staining media, permeabilized with Permeabilization buffer (eBioscience), and stained with intracellular antibodies in 500 uL for 60 min at 4°C on a shaker. Cells were washed twice in cell staining media and stained with 1mL of 1:4000 191/193Ir DNA intercalator (Fluidigm) and diluted in PBS with 1.6% paraformaldehyde overnight. Cells were washed once with cell staining media and 2 times with double-deionized water. Care was taken to assure buffers preceding analysis were not contaminated with metals in the mass range above 100 Da. Mass cytometry samples were diluted in Cell Acquisition Solution (Fluidigm) containing bead standards (see below) to approximately 10^6^ cells per mL and then analyzed on a CyTOF 2 mass cytometer (Fluidigm) equilibrated with Cell Acquisition Solution.

Data normalization was performed as previously described^68^. All mass cytometry files were normalized together using the mass cytometry data normalization algorithm^68^, which uses the intensity values of a sliding window of bead standards to correct for instrument fluctuations over time and between samples. After normalization and debarcoding of files, singlets were gated by event length and DNA. Live cells were identified by cisplatin negative cells. All positive and negative populations and antibody staining concentrations were determined by titration on positive and negative control cell populations.

Scaffold maps were generated as previously described^29^ using the open-source Statistical Scaffold R package available at github.com/SpitzerLab/statisticalScaffold. Landmark reference nodes were gated from peripheral blood mononuclear cells, while unsupervised clusters were generated via Clustering Large Applications (CLARA) clustering from samples pooled together with each sample contributing an equal number of cells. Uniform manifold approximation and projection (UMAP) was performed on ArcSinh (cofactor=5) transformed protein expression values on equal numbers of cells from each sample by randomly subsampling cells with parameters min.dist = 1.0. Clusters were identified by CLARA clustering using cells from all samples concatenated together.

### Proteomic mass spectrometry

HSC and HEI-193 media samples were prepared for mass spectrometry as described above, and 50 μL of media was mixed with 50 μL of lysis buffer containing 75 mM ammonium bicarbonate, 1% sodium deoxycholate, 5 mM TCEP and 40 mM chloroacetamide and incubated at 37°C for 30 minutes to reduce and alkylate proteins. Proteins were digested with endoproteinase LysC (Wako Chemicals) and trypsin (Promega) overnight at 37°C, followed by sodium deoxycholate and 2% TFA precipitation and centrifugation for 15 minutes at maximum speed. Peptides were desalted using MicroSpin Columns (The Nest Group) and resuspended in 4% formic acid and 3% acetonitrile. Approximately 1 μg of digested peptides per sample were loaded onto a 75 μm ID column packed with 25 cm of Reprosil C18 1.9 μm, 120 Å particles (Dr. Maisch GmbH HPLC), and eluted into an Orbitrap Fusion Lumos Tribrid mass spectrometer (Thermo Fisher Scientific) over the course of a 120-minute acquisition by gradient elution delivered by an Easy1200 nLC system (Thermo Fisher Scientific). The composition of mobile phase A and B were 0.1% formic acid in water and 0.1% formic acid in 80% acetonitrile, respectively. All MS spectra were collected with orbitrap detection and the most abundant ions were fragmented by higher energy collision dissociation and detected in the ion trap, with a 1 second cycle time between MS1 spectra. All data were searched against the Uniprot human proteome database. Peptide and protein identification searches were performed using the MaxQuant data analysis algorithm, and all peptide and protein identifications were filtered to a 1% false-discovery rate^69,70^. Label-free quantification and statistical testing was performed using the MSstats statistical R-package^71^. The mass spectrometry data files (raw and search results) have been deposited to the ProteomeXchange Consortium via the PRIDE partner repository with dataset identifier PXD014798^72^. Parallel reaction monitoring (PRM) measurements were acquired by LC-MS/MS on a Q-Exactive Plus (Thermo Fisher Scientific) mass spectrometer equipped with an EASY-nLC 1200 system (Thermo Fisher Scientific) using the same column configuration and HPLC settings as described above. All PRM data was analyzed by Skyline 4.2.0.19072^73^, quantified via MSstats, and have been deposited to Panorama Public^74^ with dataset identifier PXD014883.

### Targeted metabolite liquid chromatography mass spectrometry

HEI-193 cells were cultured and treated as described above, and 100uL of cell free supernatant (CFS) or cell pellets containing 10 million cells each were separately combined with cold H2O:MeOH at a ratio of 1:1 containing internal standards of 2-amino-3-bromo-5-methylbenzoic acid and a labeled amino acid mixture (13C,15N) spiked-in at 1ug/mL and 10nM/mL, respectively. CFS and cell pellets were briefly vortexed or rotated overnight, respectively, and immediately chilled at 4°C. All samples were centrifuged at 4°C for 30 minutes at max g and supernatant was aliquoted for quantification.

Targeted LCHRMS analysis was performed on a Sciex Exion UPLC equipped with a C18 Polar column (Phenomenex, 150 × 2.1 mm, 2.6 um) and coupled to a Sciex QTRAP 7500 operated in negative mode with MRM acquisition of MS/MS data. A gradient of 0% - 90% acetonitrile over 16 min was used for compound separation. The XICs of all samples and QCs were visually inspected. All QCs displayed appropriate signals for analysis in addition to a significant peak capacity and retention time reproducibility. Raw data files were uploaded to SciExOS for processing where filtering, feature detection, integration, and alignment of the chromatograms in each sample was performed. As a result, data matrices were generated with intensity values representing the area under the curve of significant chromatographic peaks above the lower limit of detection.

### Primary schwannoma cell culture, treatment, and metabolomic mass spectrometry

Primary human vestibular schwannomas were cultured as previously described^75^. Briefly, tumors were collected from the operating room and placed on ice for transfer to the lab. Under sterile conditions in a cell culture hood, the tissue was minced into 1 mm^3^ pieces then digested for 45 minutes at 37°C in 0.25% trypsin/0.1% collagenase solution. Cells were resuspended in Dulbecco’s Modified Eagle Medium supplemented with insulin (10 µg/mL, Sigma), 10% fetal bovine serum, and N2 supplement (Sigma). The cell suspension was plated onto culture dishes pre-treated with poly-L-ornithine and laminin. Cultures were placed in a humidified incubator at 37°C with 5.0% CO2 and grown for 7-14 days without passage. Media was exchanged every 2-3 days. Cultures were treated with 0, 3, 10, or 20Gy gamma-irradiation from a radioactive Cesium source. 6 or 72 hours after radiotherapy, cultures were washed with ice cold PBS and water, and flash-frozen in liquid nitrogen. Derivatized metabolite extracts from these primary schwannoma cell culture were analyzed by Gas Chromatography-Mass Spectrometry on a ThermoISQ Quadrupole. Raw data were analyzed with TraceFinder 4.1 (Thermo). Cell culture data were normalized to total ion signal to control for extraction, derivatization, and/or loading effects.

### Quantitative polymerase chain reaction

RNA was isolated using the RNEasy Mini Kit and a QiaCube (QIAGEN), and cDNA was synthesized using the iScript cDNA Synthesis Kit (Bio-Rad, Hercules, CA) and a ProFlex thermocycler (Thermo Fisher Scientific). Target genes were amplified using PowerUp SYBR Green Master Mix and a QuantStudio 6 thermocycler (Thermo Fisher Scientific). Gene expression was calculated using the ΔΔCt method for candidate genes, with normalization to *GAPDH* (Supplementary Table 11).

### Bulk RNA sequencing and analysis

RNA was isolated from frozen tumors or cell cultures using the All-Prep Universal Kit, and clean-up was performed using the RNEasy Kit as needed (Qiagen). RNA sequencing libraries were generated using the Illumina TruSeq Stranded mRNA Library Prep Kit and sequenced on an Illumina HiSeq-4000 using the paired-end 100 protocol for schwannomas and single-end 50 protocol for HEI-193 schwannoma cells. Reads were aligned to GRCh38 using the splice-aware aligner HISAT2 version 2.0.3 against an index containing SNP and transcript information (genome_snp_tran)^76^. Ensembl build 75 genes were quantified with featureCounts using uniquely mapped reads^77^. Differential expression analysis was performed using DESeq2 using the Wald test with an adjusted p-value threshold of 0.05 corrected for multiple hypotheses^78^. Complete linkage hierarchical clustering was performed using 1-Pearson correlation as the distance matrix with differentially expressed genes.

### Single-cell RNA sequencing and analysis

HEI-193 schwannoma cells were dissociated in Trypsin-EDTA 0.25% (Thermo Fisher Scientific), passed through a 40 μm strainer, and washed/resuspended in phosphate buffered saline. Fresh schwannomas were minced with sterile Bard-Parker #10 surgical scalpels (Aspen Surgical) and incubated in 0.4% Collagenase Type 2 (Worthington) in pre-oxygenated Dulbecco’s Modified Eagle Medium (Thermo Fisher Scientific) for 75 minutes at 37°C while rotating at 800 rotations per minute on a thermomixer. The suspension was sequentially filtered through 70 μm and 40 μm strainers (Thermo Fisher Scientific), centrifuged at 300g for 5 minutes, and resuspended in cold phosphate buffered saline. Single cell suspensions were loaded onto a 10x Chromium controller using the Chromium Single Cell 3’ Library & Gel Bead Kit v3 (10x Genomics). Libraries were sequenced on an Illumina NovaSeq (10X specific protocol) with >50,000 reads per cell.

Library demultiplexing, read alignment to human genome GRCh38 and UMI quantification was performed in Cell Ranger version 1.3.1 (10X Genomics). Schwannoma cells with greater than 200 unique genes detected and fewer than 15% of reads attributed to mitochondrial transcripts were retained. Data were normalized and variance stabilized by SCTransform in Seurat version 3.0 using unique molecular identifier (UMI) count and percent of reads aligned to mitochondrial transcripts as covariates^79^. UMAP was performed on significant (p<0.05) principal components (determined by JackStraw analysis) and batch corrected using Harmony with parameters min.dist=0.3. Louvain clustering was performed with resolution=0.3, and cluster markers were identified based on expression in at least 25% of cells and differentially expressed by more than 25% compared to all other clusters. For HEI-193 schwannoma cells, analysis was performed as above, except that cells with greater than 2000 UMIs and fewer than 20% of reads attributed to mitochondrial transcripts were retained, and UMAP was performed with min.dist=0.2 without harmonization. Gene ontology analysis was performed using Enrichr without inclusion of ribosomal subunits or mitochondrial genes^58^. Cell cluster identification was further analyzed using the AddModuleScore function within Seurat 3.0 to score expression signatures based on gene sets from the Molecular Signatures Database (MSigDB)^80^, intersected with differentially expressed cluster markers described above, and then normalized across all populations in UMAP space^79^.

### Single-nuclei RNA sequencing and analysis

Flash frozen archived schwannoma specimens were minced with sterile Bard-Parker #10 surgical scalpels (Aspen Surgical) and mechanically dissociated with a Pestle Tissue Grinder (size A, Thomas Scientific) in ice cold lysis buffer consisting of 0.32 M sucrose, 5 mM CaCl2, 3 mM MgAc2, 0.1 mM EDTA, 10 mM Tris-HCl, 1 mM DTT and 0.1% Triton X-100 in DEPC-treated water^81^. Sucrose solution consisting of 1.8 M sucrose, 3 mM MgAc2, 1 mM DTT and 10 mM Tris-HCl in DEPC-treated water was added to the bottom of the lysis solution in ultracentrifuge tubes (Beckman Coulter) to form a gradient, which was ultracentrifuged at 107,000g for 2.5 hours at 4°C. Nuclei pellets were resuspended in phosphate buffered saline and sequentially filtered twice in 30 μm strainers (Miltenyi Biotec). Isolated nuclei were assessed with DAPI staining and loaded onto a 10x Chromium controller using the Chromium Single Cell 3’ Library & Gel Bead Kit v2 (10x Genomics). Library sequencing and preprocessing for single nuclei was performed as described above for single-cell libraries, except that a pre-mRNA reference library (GRCh38) including intronic segments was used for read alignment and quantification. Nuclei libraries with greater than 400 unique genes detected and fewer than 5% of reads attributed to mitochondrial transcripts were retained.

To integrate single-nuclei and single-cell RNA sequencing data, all libraries passing respective QC filters described above were combined *in silico* and normalized with variance stabilization using SCTransform in Seurat version 3.0, with UMI count, percent of reads aligned to mitochondrial transcripts, and technique (single-cell versus single-nuclei) as covariates^79^. Principal component analysis was performed, and single nuclei and single cell data were harmonized using Harmony in Seurat version 3.0 using technique and day-of tumor isolation as covariates. Clustering and marker identification was performed as described above for single cell-only analysis, with parameters min.dist=0.3 and resolution=0.3.

### CRISPR interference genome wide screening

CRISPRi screens were performed as described previously^65,82^. Briefly, HEI-193 cells stably expressing CRISPRi components (dCas9-KRAB) were transduced with lentivirus supernatant containing the third generation dual sgRNA CRISPRi library, which targets 20528 genes and 1025 sgNTC^82^. Screens were performed in triplicate cultures with coverage of at least 500x cells per construct. sgRNA expressing cells were selected using puromycin (1 µm/mL) for 48 hours and transferred to puromycin-free normal growth media for 48 hours to allow recovery. Initial (T0) cell populations were then frozen in 10% DMSO and processed for genomic DNA alongside endpoint (T12) cell populations, which were obtained after 10 cell doublings. Triplicate screens were also performed with radiotherapy (1.8Gy x 5 fractions) delivered daily starting on T0. Genomic DNA was harvested using the NucleoSpin Blood L Kit (Machery-Nagel) for each cell population, and sgRNA cassettes were amplified using 22 cycles of PCR using NEBNext Ultra II Q5 PCR MasterMix (New England Biolabs). Sequencing was performed on a NovaSeq 6000 (Illumina) using custom sequencing primers^82^.

sgRNA read counts were aligned using custom Python scripts without allowing mismatches. sgRNA counts with discordant target genes from the same vector, representative of vector recombination, or fewer than 100 reads detected in the T0 populations, were discarded from downstream analysis. Growth phenotype (gamma) was defined as log2(sgRNA count T12 / sgRNA count T0) minus median sgNTC log2(sgRNA count T12 / sgRNA count T0) as previously described^65^. Radiation phenotype (rho) was defined as log2(sgRNA count T12 (1.8Gy x 5) / sgRNA count T12 (0 Gy)). Statistical significance was quantified using a two-sided Student’s t-test comparing replicate distributions of library-normalized counts for each sgRNA between conditions (rho) or time points (gamma). A discriminant threshold of 5, derived from the product of normalized gene phenotype and -log10(p-value), corresponding to an empiric false discovery rate of ∼1%, was selected for hit definitions^65^.

### Perturb-seq

The single-cell Perturb-seq library was composed of sgRNAs targeting genes from RNA sequencing and DNA methylation profiling of human vestibular schwannomas (Supplementary Table 11). Library cloning was performed as previously described^83^. Briefly, insert lyophilized oligonucleotides (Twist Biosciences) containing the sgRNA sequences were resuspended at 100 nM in H_2_O and amplified by PCR using HF Phusion polymerase (New England Biolabs). After verifying amplification on a 10% acrylamide gel, the PCR product was purified using MinElute Cleanup Kit (Qiagen). The pBA904 Perturb-seq sgRNA vector backbone (Addgene, #122238) was digested using FastDigest BstX1 and Blp1 (ThermoFisher Scientific) and excised from a 0.8% agarose gel for purification using the NucleoSpin Gel Cleanup Kit (Macherey-Nagel). The insert pool and digested backbone were then ligated overnight at 16°C using T4 DNA Ligase (New England Biolabs) with a vector-to-insert ratio of 1:1. The ligation reaction was purified by ethanol precipitation and transformed into Stellar Competent Cells (Clontech) to assess library diversity by Sanger sequencing of 10 clones. For large-scale transformation, the library was transformed into MegaX Electrocompetent cells (ThermoFisher Scientific) using electroporation on 15-cm LB carbenicillin plates. The library stock was prepared by scraping plates followed by purification using the Midi Prep kit (Macherey-Nagel). Library sequencing was performed on a MiSeq run to ensure sgRNA uniformity. Using lentivirus, cells were transduced with the library at 1,000X coverage at an MOI of 0.1, corresponding to approximately 95% of cells with a single integration. 72 hours following transduction, sgRNA+ cells were FACS sorted (BFP+) and were recovered in DMEM 10% FBS. Sorted cultures were treated with either 0Gy, 1.8Gy x 5 daily fractions, or 12.5Gy x 1 fraction of radiotherapy using the X-Rad 320 irradiator (Precision X-Ray). 12 hours following the final fraction of radiotherapy, cells were trypsinized and harvested in single cell suspension on the 10x Chromium Controller (10x Genomics).

Single-cell Perturb-seq libraries were processed using the Chromium Next GEM Single Cell 3’ GEM, Library & Gel Bead Kit v3.1with Feature Barcoding (10x Genomics), allowing direct capture of modified sgRNAs^83^, and sequenced on a illumina NovaSeq-6000. Cells from the 0Gy condition were run across 2 GEM groups, while each of the radiation conditions were run on its own GEM group. Library demultiplexing, gene expression read alignment to human genome GRCh38, UMI quantification, and sgRNA assignment and quantification were performed in Cell Ranger version 6.1.2 with sgRNA barcoding (10X Genomics).

Data analysis was performed using Seurat 4.3.0 in R version 4.2.2. Cells with greater than 200 detected features and UMIs mapping to only one sgRNA were retained. Target gene knockdown was quantified by library normalizing the transcriptome of each cell and obtaining the mean expression of each gene across all cells belonging to a given sgRNA within an individual GEM group (pseudobulk). RNA remaining for each gene target was calculated by dividing the pseudobulk expression in on-target cells with those of sgNTC cells, with a pseudocount of 0.01 added to each component. Only sgRNAs exerting greater than 75% knockdown in at least one of the two 0 Gy conditions, and also represented in at least 10 cells in each condition, were retained for analysis. To score expression changes of gene modules, marker genes were derived from Louvain clustering of human schwannoma cell types (Fig. 1b) and HEI-193 schwannoma cell states (Fig. 2c), as described above. The top 10 most specific cluster markers were used to interrogate each cell type or cell state, and the mean pseudobulk expression of these markers were obtained for each on-target sgRNA as well as for sgNTC cells, in each radiotherapy condition. sgRNA phenotypes were then normalized to pseudobulk cells containing sgNTC in the 0 Gy condition. These fold changes were log_2_ transformed and grouped within radiotherapy conditions using hierarchical clustering with Pearson correlation and complete linkage.

Differential expression analysis in Perturb-seq data was performed using the FindMarkers function in Seurat with MAST as the test type. Genes with p<0.05 and |log_2_(fold change)|>1) were considered differentially expressed and used for gene ontology analyses in EnrichR with separate queries for all enriched or suppressed genes. To identify perturbations which preferentially generated phenotypes in radiotherapy conditions, the number of significantly differentially expressed genes were compared to identify perturbations with more than 40 excess differentially expressed genes in the radiotherapy (1.8Gy x 5) conditions compared to control (0Gy) conditions.

### Single-nuclei ATAC, RNA, and CRISPRi perturbation sequencing (snARC-seq)

Genetic perturbations in snARC-seq reply on the CROP-seq vector (Addgene, pBA950)^84^, which allows for capture of sgRNA identity from nuclear RNA transcripts. Oligonucleotides containing protospacer sequences (Supplementary Table 11) were ordered as a pool from Twist Bioscience and PCR amplified as described above. The oligonucleotide pool was ligated into the Crop-seq backbone using BstXI and BlpI digestion and T4 ligation. Library representation, including sgNTC overrepresentation, was performed on a MiSeq run to ensure sgRNA uniformity.

The cloned library was packaged into lentivirus using HEK-293T cells. HEI-193 cultures were transduced to an MOI of 0.1 and FACS sorted for sgRNA+ cells after 48 hours. Radiotherapy was then delivered to either 0Gy or 1.8Gy x 5 daily fractions using the X-Rad 320 irradiator (Precision X-Ray). Following completion of radiotherapy, cells were harvested with Trypsin and prepared following the Nuclei Isolation for Single Cell Multiome ATAC + Gene Expression Sequencing 10x Protocol (10x Genomics, CG000365 Rev C). Briefly, 1-2 10^5^ cells were incubated in chilled Lysis Buffer (Tris-HCl (pH 7.4) 10 mM, NaCl 10 mM, MgCl2 3 mM, Tween-20 0.1%, Nonidet P40 Substitute 0.1%, Digitonin 0.01%, BSA 1%, DTT 1mM, RNase inhibitor 1 U/µL) on ice for 4 minutes, washed with Wash Buffer (Tris-HCl (pH 7.4) 10 mM, NaCl 10 mM, MgCl2 3 mM, Tween-20 0.1%, BSA 1%, DTT 1 mM, RNase inhibitor 1 U/uL) 3 times, and resuspended in diluted nuclei buffer. Nuclei were counted and membrane integrity was evaluated by Trypan staining on the Countess II FL Automated Cell Counter (Thermo). Nuclei suspensions were diluted to approximately 3,000 nuclei/µL and processed according to the 10x Chromium Next GEM Single Cell Multiome ATAC + Gene Expression protocol (CG000338 Rev A).

To recover sgRNA identities from single-nuclei RNA fractions, CROP-seq guides were amplified into dual-indexed Illumina libraries from the cDNA product of the 10x multiome protocol as described above with a three-round hemi-nested PCR as previously described^85^. 15 ng of full-length cDNA product was amplified with primers binding to the sgRNA constant region and 10x Genomics Read 1 Adaptor. Two subsequent PCR steps were performed to introduce i5 and i7 indices and Illumina P5 and P7 adaptors. After the first round of PCR, each product was size-selected using SPRI beads at 1.0x. Subsequent PCRs were conducted with 1 ng product. The final PCR product was size selected with 0.5x and then 1.0x SPRI beads and library quality was assessed by TapeStation high-sensitivity D1000 analysis (Thermo). CROP-seq libraries were pooled with gene expression libraries and sequenced on an Illumina NovaSeq 6000 using the paired end 100 bp protocol.

Library demultiplexing, read alignment to human genome GRCh38, and UMI quantification for the RNA and ATAC fractions were performed using Cell Ranger ARC version 2.0.1 (10X Genomics). Crop-seq sgRNAs were detected using kallisto bustools^86^. First, a sgRNA reference was built using kb ref which was used as a reference for kb count including the ARC multiome Gene Expression Whitelist version 1 (10x Genomics). For each cell barcode group, sgRNAs with less than 6 UMIs were filtered and sgRNAs were assigned to cells with a hypergeometric test using Geomux. The resulting sgRNA assignments were used to determine cell barcode groups with a single detected sgRNA.

Preprocessing of single-nuclei RNA and ATAC data was performed using Signac v1.8.0^87^. Cells containing the following quality measures were retained: ATAC UMI between 1000 and 100000, RNA UMI between 50 and 25000, nucleosome signal < 4, and TSS enrichment > 1. ATAC peak calling was performed using the MACS2 wrapper in Signac. Peaks were processed using term frequency inverse document frequency (TF-IDF) normalization and singular value decomposition. RNA counts were normalized and variance stabilized by SCTransform using default parameters. ATAC UMAP projection was calculated using the latent semantic indexing reduction dimensions 2 to 30. RNA UMAP projection was calculated using principal component analysis dimensions 1:20, which was empirically determined using ElbowPlot. Louvain clustering for either ATAC or RNA data was performed using resolution parameter of 0.5.

Gene activity scores from ATAC signal was generated by quantifying ATAC UMIs mapped to the promoter of each gene, defined as between the TSS and 2000 bp upstream of the TSS for each gene. To score gene activity changes of gene modules between radiotherapy and control conditions, gene activity score were caculated for marker genes derived from Louvain clustering of human schwannoma cell types (Fig. 1b) and HEI-193 schwannoma cell states (Fig. 2c), as described above. The top 10 most specific cluster markers were used for each cell type or cell state, and the mean pseudobulk gene activity of these markers were obtained for each on-target sgRNA as well as for sgNTC cells, in each radiotherapy condition. sgRNA phenotypes were then log_2_ transformed and the gene activity vector in radiotherapy (1.8Gy x 5) conditions were subtracted by those in control conditions (0Gy). These fold changes were clustered using hierarchical clustering with Pearson correlation and complete linkage.

Transcription factor motif deviations in the setting of genetic or therapeutic perturbations were quantified using the ChromVAR^50^ wrapper in Signac with default parameters against the peaks assay. Mean motif deviations for each sgRNA identity in each condition were quantified and subsetted to only motifs whose cognate transcription factors were expressed ion HEI-193 cells using RNA sequencing data, as described above. To estimate the differential motif deviations for a given sgRNA perturbation in radiotherapy conditions versus control conditions, deviations were log_2_ transformed and deviations in radiotherapy conditions were subtracted by those in control conditions. To further quantify chromatin accessibility at motif regions as consequences of epigenetic regulator snARC-seq perturbations, differentially accessible regions were determined using the FindMarkers function in Signac using logistic regression. Differentially accessible regions were further subsetted by those whose nearest neighbor gene was associated with ChIP-seq peaks derived from the ENCODE project (KLF13 accession ENCFF453MMH, SETDB1 accession ENCFF773RNU). Average ATAC profiles were obtained for each sgRNA condition at these restricted sets of differentially accessible regions with a window *±* 1000 bp and then normalized to the average read depth at the 100 bps at each end flanks of the distributions.

### Statistics

All experiments were performed with independent biological replicates and repeated, and statistics were derived from biological replicates. Biological replicates are indicated in each figure panel or figure legend. No statistical methods were used to predetermine sample sizes, but sample sizes in this study are similar or larger to those reported in previous publications. Data distribution was assumed to be normal, but this was not formally tested. Investigators were blinded to conditions during clinical data collection and analysis of mechanistic or functional studies. Bioinformatic analyses were performed blind to clinical features, outcomes or molecular characteristics. The clinical samples used in this study were retrospective and nonrandomized with no intervention, and all samples were interrogated equally. Thus, controlling for covariates among clinical samples is not relevant. Cells were randomized to experimental conditions. No clinical, molecular, or cellular data points were excluded from the analyses. Unless specified otherwise, lines represent means, and error bars represent standard error of the means. In general, statistical significance is shown by asterisks (*p<0.05, **p*≤*0.01, ***p*≤*0.0001), but exact p*-*values are provided in the figure legends when possible. Lines show means ± standard error of the means. Boxplots show 1^st^ quartile, median and 3^rd^ quartile; whiskers represent 1.5 inter-quartile range, and data outside this range are shown by points. Results were compared using Student’s *t*-tests, Fisher’s exact tests, and ANOVA, which are indicated in figure legends alongside approaches used to adjust for multiple comparisons. Multiple linear regression was performed in *R* (v 3.6.0). Recursive portioning analysis was performed using the *rpart* package in *R*, with minimum split size of 10 and complexity parameter of 0.02. Variables with p-value of 0.1 or less on univariate analysis were included in RPA and multivariate analysis. Logistic regression models were constructed and evaluated using receiver operating characteristic analysis. A nomogram was created using the *rms* package in R, which generates a graphical nomogram based on a general linear model.

## Figures

**Extended Data Fig. 1.**
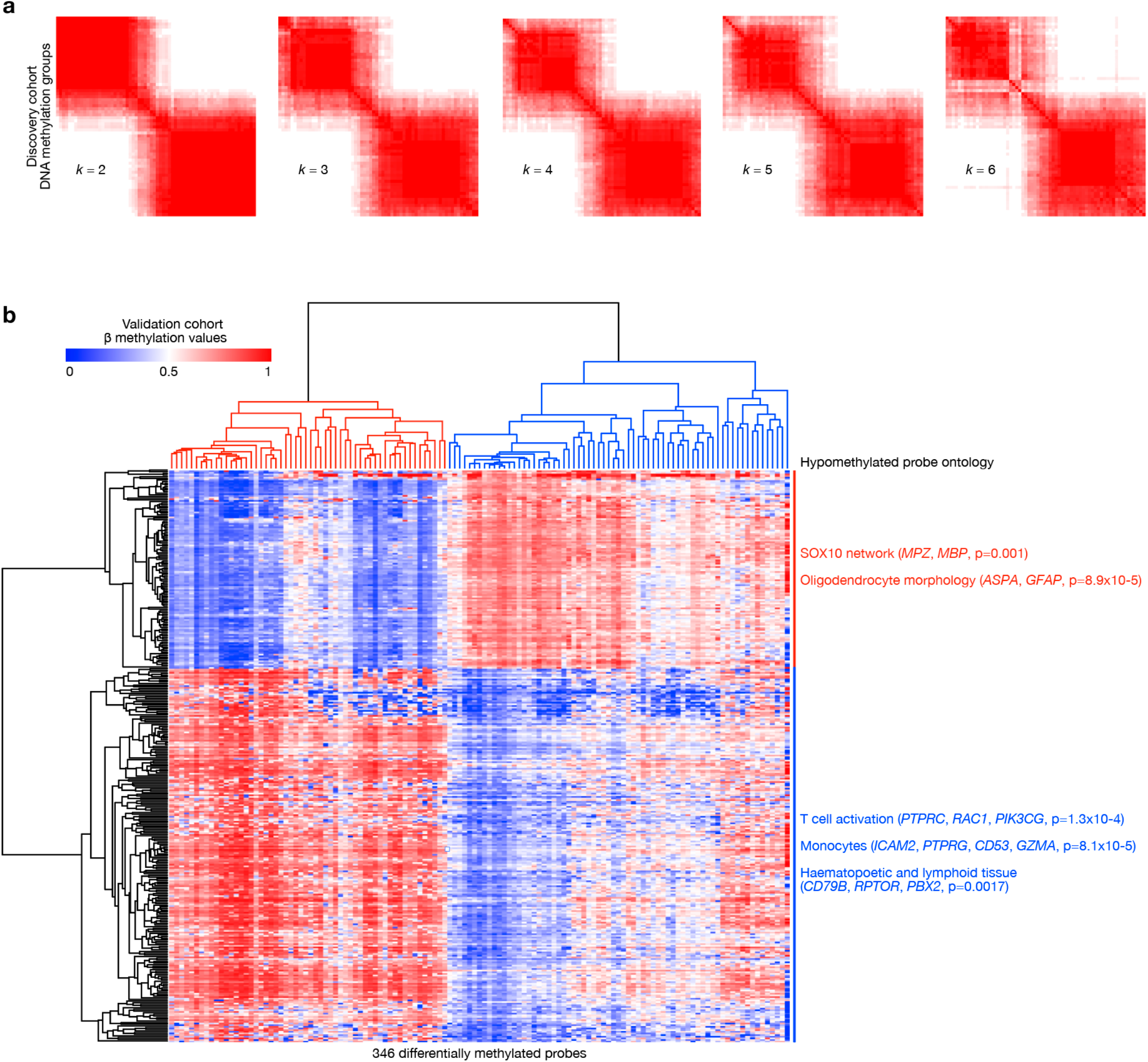
DNA methylation-based classification of schwannomas identifies 2 molecular groups. **a**, Consensus clustering of pairwise Pearson correlation coefficients from 850k DNA methylation profiling of 66 vestibular schwannomas (k=2-6) comprising the discovery cohort from UCSF. **b**, Hierarchical clustering of 450K DNA methylation profiles from 125 vestibular or spinal schwannomas comprising the validation cohort^14^ using DNA methylation probes overlapping with the top 2000 differentially methylated probes from the discovery cohort (346 probes, Fig. 1a). Significant gene ontology terms of hypomethylated probes distinguishing molecular groups are shown, validating neural crest schwannomas (NCS) and immune-enriched schwannomas (IES).

**Extended Data Fig. 2.**
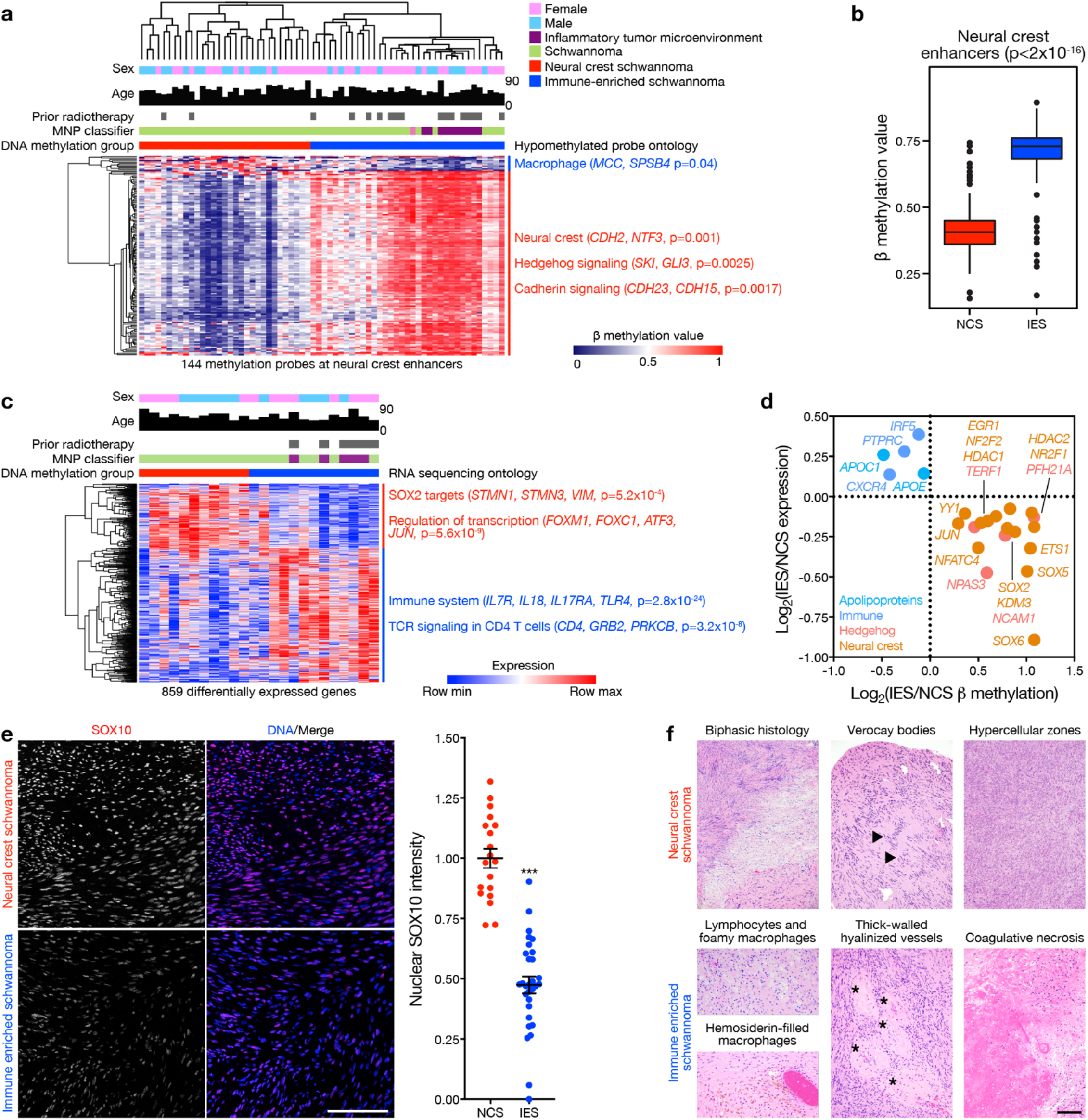
Epigenetic, transcriptomic, protein expression, and histologic validation of neural crest and immune enriched schwannomas. **a**, Hierarchical clustering of schwannomas (n=66) according to DNA methylation probes overlapping with neural crest enhancers. Significant gene ontology terms corresponding to hypomethylated probes in neural crest schwannomas (NCS) or immune-enriched schwannomas (IES), clinical metadata, and the molecular neuropathology (MNP) DNA methylation classification of central nervous system tumors^18^ are shown. **b**, Distribution of median schwannoma β methylation values for DNA methylation probes overlapping with neural crest enhancers. P-value determined using two-sided Kolmogorov-Smirnov test. Boxplots show 1^st^ quartile, median, and 3^rd^ quartile. Whiskers represent 1.5 inter-quartile range, and data outside range are shown. **c**, Hierarchical clustering of differentially expressed genes from RNA sequencing of NCS or IES (n=24). Significant gene ontology terms corresponding to enriched genes in each molecular group and meta-data are shown as in **a. d**, Select schwannoma apolipoprotein, immune, Hedgehog, and neural crest genes with anti-correlated expression and β methylation from 24 tumors with matched RNA sequencing and DNA methylation profiling. **e**, Quantitative immunofluorescence microscopy for the schwannoma differentiation marker SOX10 across NCS or IES (n=49). Scale bar, 100μm. Lines represent means and error bars represent standard error of means (Student’s t test, ***p≤0.0001). **f**, H&E-stained sections of schwannomas (n=66) showing biphasic histology (90% versus 67%, p=0.0001), Verocay bodies (arrows, 84% vs 47%, p<0.0001) and hypercellular zones (30% vs 3%, p=0.0001) in NCS versus lymphocyte and macrophage infiltration (59% versus 17%, p<0.0001), hemosiderin deposition (93% versus 68%, p<0.0001), hyalinized or thick-walled vessels (asterisks, 74% versus 57%, p=0.017), and necrosis (18% versus 3%, p=0.0008) in IES. Fisher’s exact tests. Scale bar, 100μm.

**Extended Data Fig. 3.**
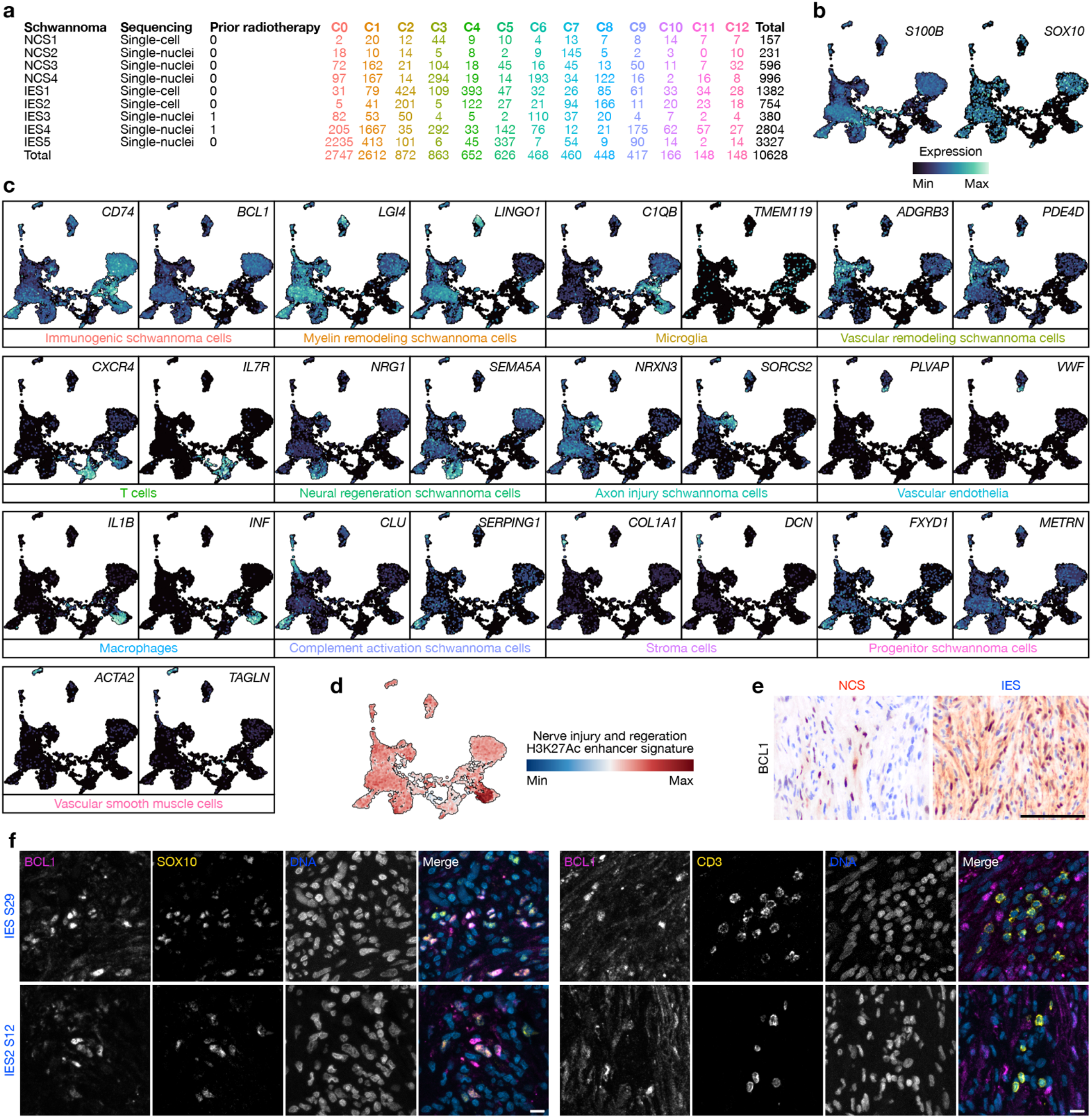
Integrated schwannoma single-nuclei and single-cell RNA sequencing. **a**, Distribution of integrated, harmonized schwannoma single-nuclei (n=6) and single-cell RNA sequencing (n=3) transcriptomes (Fig. 1b). **b**, Feature plot of integrated UMAP from harmonized schwannoma single-nuclei and single-cell RNA sequencing showing expression of the schwannoma differentiation marker genes *S100B* (left) or *SOX10* (right). **c**, Feature plots of schwannoma and microenvironment cell type marker genes. Immunogenic schwannoma cells were distinguished by enrichment of *CD74*, which mediates macrophage migration, axon repair, and survival of neural progenitor cells^23,24^. Myelin remodeling schwannoma cells were distinguished by enrichment of *LINGO1*, which regulates myelin recovery in demyelinating disease and contributes to nerve regeneration^88,89^. Vascular remodeling schwannoma cells were distinguished by enrichment of *ADGRB3*, an angiogenesis regulator that contributes to cell proliferation through the tumor microenvironment^25–27^, and the balance of angiogenesis factors is a hallmark of schwannoma^90^. Neural regeneration schwannoma cells were distinguished by enrichment of *SEMA5A*, an inhibitory axon guidance molecule that facilitates schwannoma cell proliferation by modulating axon growth^91,92^. Axon injury schwannoma cells were distinguished by enrichment of *SORCS2*, which is induced in peripheral nerve injury^93^, and *NRXN3*, which localizes to injured axons^28^. Complement pathway activation schwannoma cells were distinguished by enrichment of *CLU* and *SERPING1*, which are activated in both ischemic and traumatic brain injury^94–96^. Progenitor schwannoma cells were distinguished by enrichment of *FXYD1* and *METRN*, which are expressed in adult neural stem cells^97^. **d**, Mean expression signature of genes neighboring enhancers that are activated during nerve injury and regeneration in integrated schwannoma transcriptomes^21^. **e**, IHC for the immunogenic schwannoma cell marker BCL1 reveals enrichment in IES compared to NCS (Fisher’s exact test, p=0.05). Scale bar, 100μm. **f**, Co-immunofluorescence microscopy for the immunogenic schwannoma cell marker BCL1 with the T cell marker CD3 (right) or the schwannoma differentiation marker SOX10 (left) in IES, demonstrating BCL1 expression from schwannoma cells but not from lymphocytes, in support of single-nuclei and single-cell RNA sequencing data (Extended Data Fig. 3c). Scale bar, 10μm.

**Extended Data Fig. 4.**
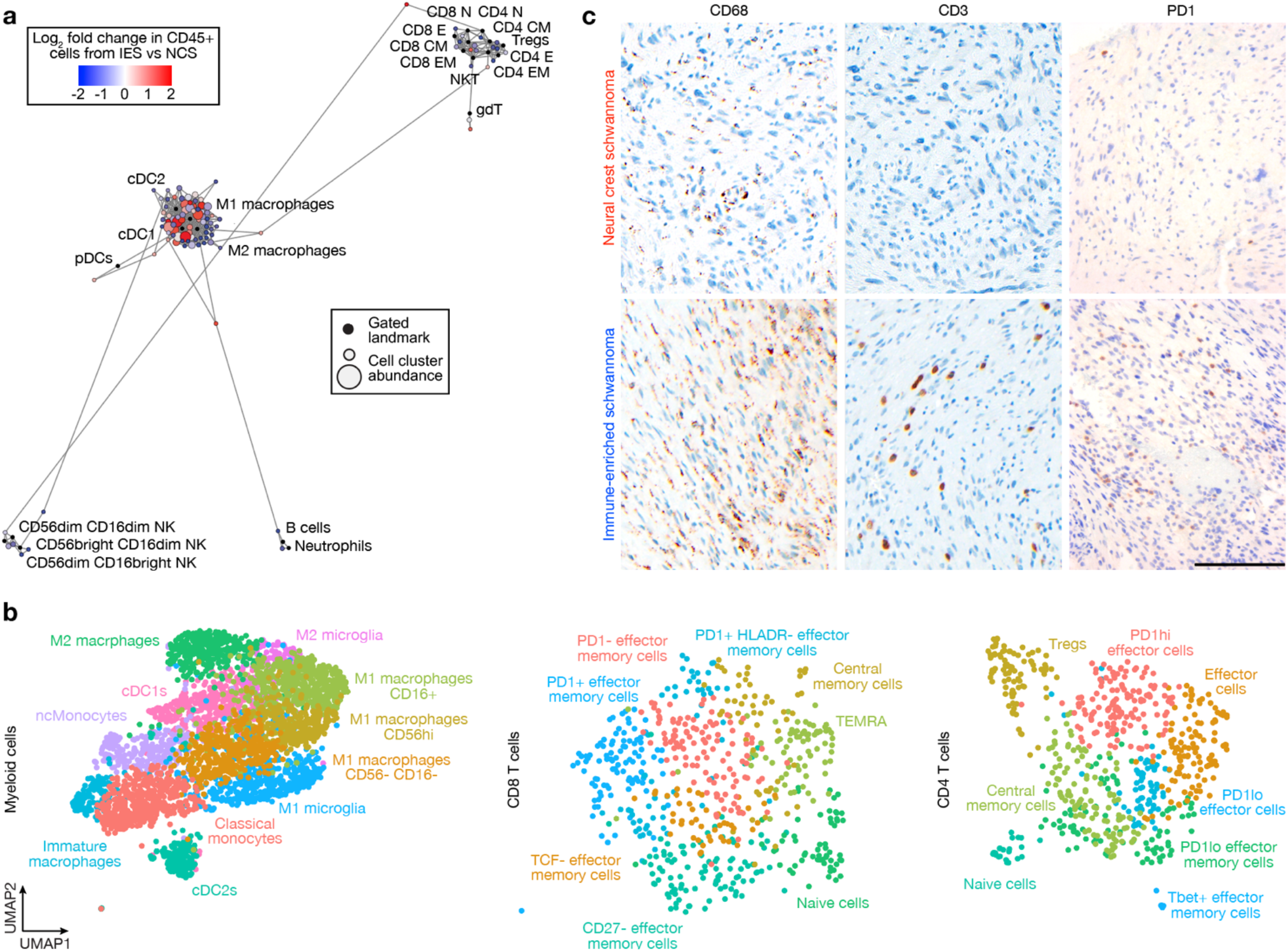
Neural crest and immune enriched schwannomas are distinguished by myeloid and CD8 T cell populations. **a**, Scaffold plot comprised of 375,355 immune cells from NCS (n=3) or IES (n=3) analyzed using mass cytometry time-of-flight (CyTOF). Manually gated landmark immune cell populations (black) are annotated. **b**, UMAP representation of myeloid cells, CD8 T cells, and CD4 T cells from schwannomas analyzed using CyTOF. **c**, Immunohistochemistry staining for CD68 macrophages (p=0.05), CD3 T cells (p=0.04), and PD1 T cells (p=0.008) reveals enrichment in IES relative to NCS (n=66). Fisher’s exact tests. Scale bar, 100μm.

**Extended Data Fig. 5.**
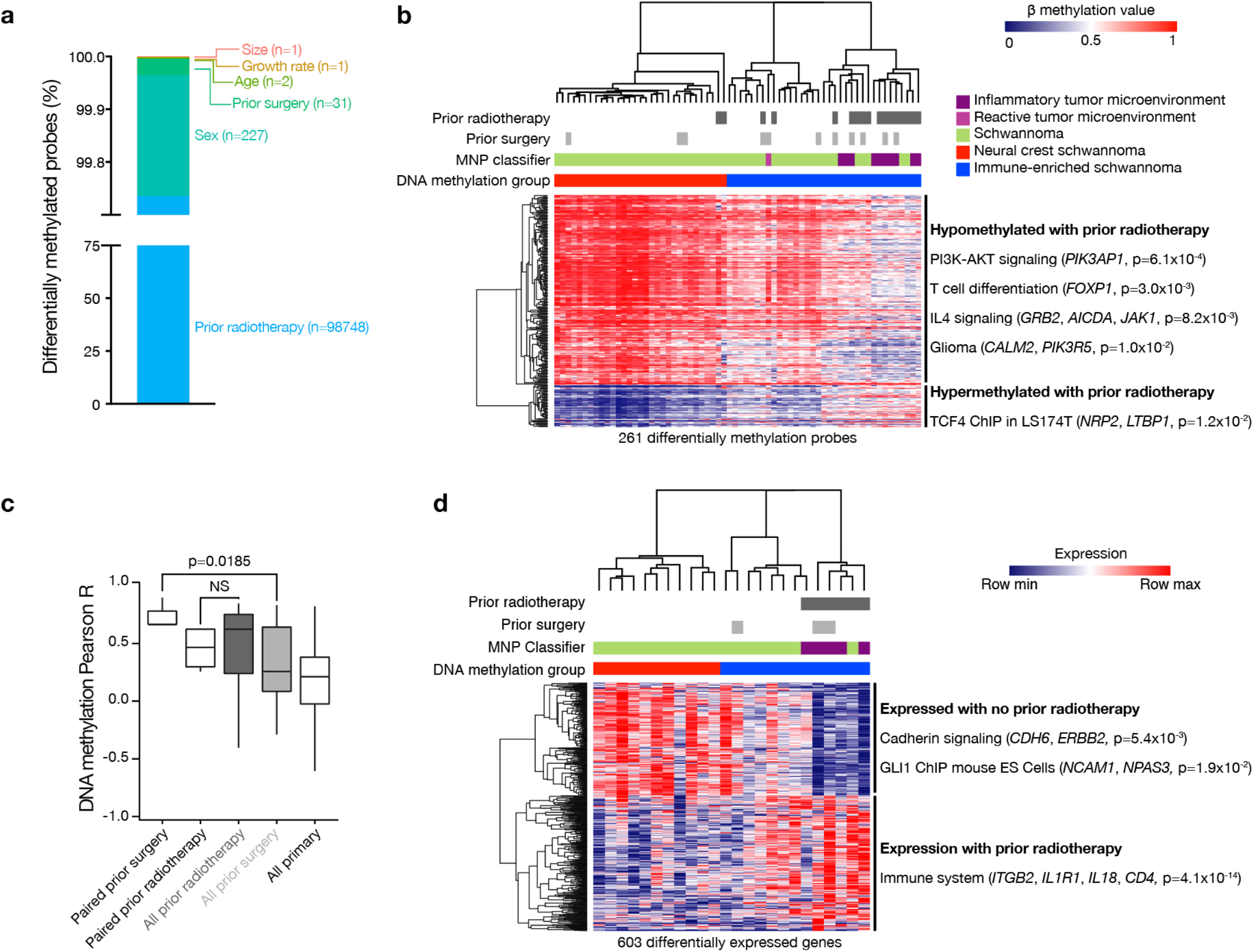
Radiotherapy is associated with epigenetic reprogramming of neural crest to immune-enriched schwannoma. **a**, Differentially methylated DNA probes based on clinical characteristics of schwannoma patients (n=66, FDR<0.05). **b**, Hierarchical clustering of differentially methylated DNA probes from schwannomas (n=66) with versus without prior radiotherapy (FDR<0.01). Significant gene ontology terms corresponding to hypomethylated probes, clinical metadata, and the molecular neuropathology (MNP) DNA methylation classification of central nervous system tumors^18^ are shown. **c**, Distributions of pairwise Pearson correlation coefficients for patient-matched pairs of primary and recurrent schwannomas. P-values determined using two-sided Kolmogorov-Smirnov test. Boxplots show 1^st^ quartile, median, and 3^rd^ quartile. Whiskers represent 1.5 inter-quartile range. **d**, Hierarchical clustering of differentially expressed genes from RNA sequencing of schwannomas with versus without prior radiotherapy (n=24). Significant gene ontology terms *±* prior radiotherapy and meta-data are shown as in **b**.

**Extended Data Fig. 6.**
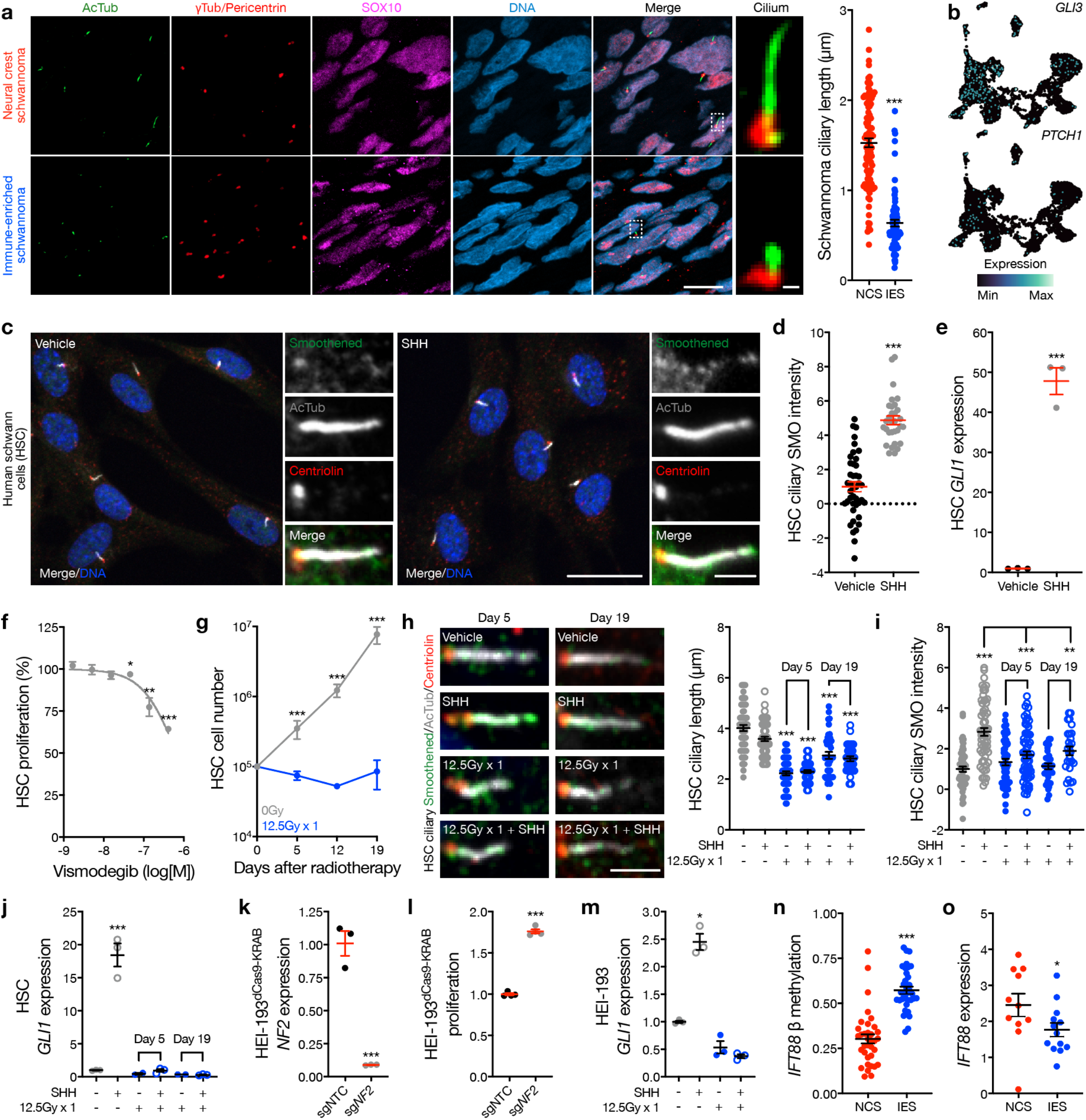
Radiotherapy inhibits Schwann and schwannoma cell Hedgehog signaling. **a**, Quantitative immunofluorescence microscopy for the ciliary axoneme marker acetylated tubulin (AcTub), the ciliary base markers gamma tubulin (γtub) and pericentrin (red), and the schwannoma cell marker SOX10 (purple) in 174 cilia from 6 schwannomas. Scale bars, 10μm and 1μm. **b**, Feature plot of integrated UMAP from harmonized schwannoma single-nuclei and single-cell RNA sequencing (Fig. 1b) showing expression of the Hedgehog target genes *GLI3* or *PTCH1* in schwannoma cells. **c** and **d**, Quantitative immunofluorescence microscopy for AcTub, the ciliary base marker Centriolin, and the Hedgehog pathway activator Smoothened in immortalized human Schwann cells (HSCs) treated with recombinant sonic Hedgehog (SHH) or vehicle control. Scale bars, 20μm and 2μm. **e**, QPCR for the Hedgehog target gene *GLI1* in HSC cultures treated with SHH or vehicle control. **f**, Quantification of HSC proliferation after treatment with the Smoothened antagonist vismodegib for 72h. **g**, HSC proliferation after treatment with radiotherapy compared to control. **h** and **i**, Quantitative immunofluorescence imaging of primary cilia in HSC cultures 5 days or 19 days after treatment with radiotherapy or control, and SHH or vehicle control. **j**, GLI1 QPCR in HSC cultures 5 days or 19 days after treatment with radiotherapy or control, and SHH or vehicle control. **k**, QPCR validation of *NF2* CRISPRi suppression in HEI-193 schwannoma cells compared to non-targeting control sgRNAs (sgNTC). **l**, Quantification of HEI-193 cell proliferation following CRISPRi suppression of *NF2* compared to sgNTC. **m**, GLI1 QPCR in HEI-193 cells 5 days after treatment with radiotherapy or control plus SHH or vehicle control. **n**, DNA methylation β values for the ciliary axoneme gene *IFT88* in NCS compared to IES. **o**, RNA sequencing transcripts per million expression of *IFT88* in NCS compared to IES. Lines represent means and error bars represent standard error of means (Student’s t tests, *p≤0.05, **p≤0.01, ***p≤0.0001).

**Extended Data Fig. 7.**
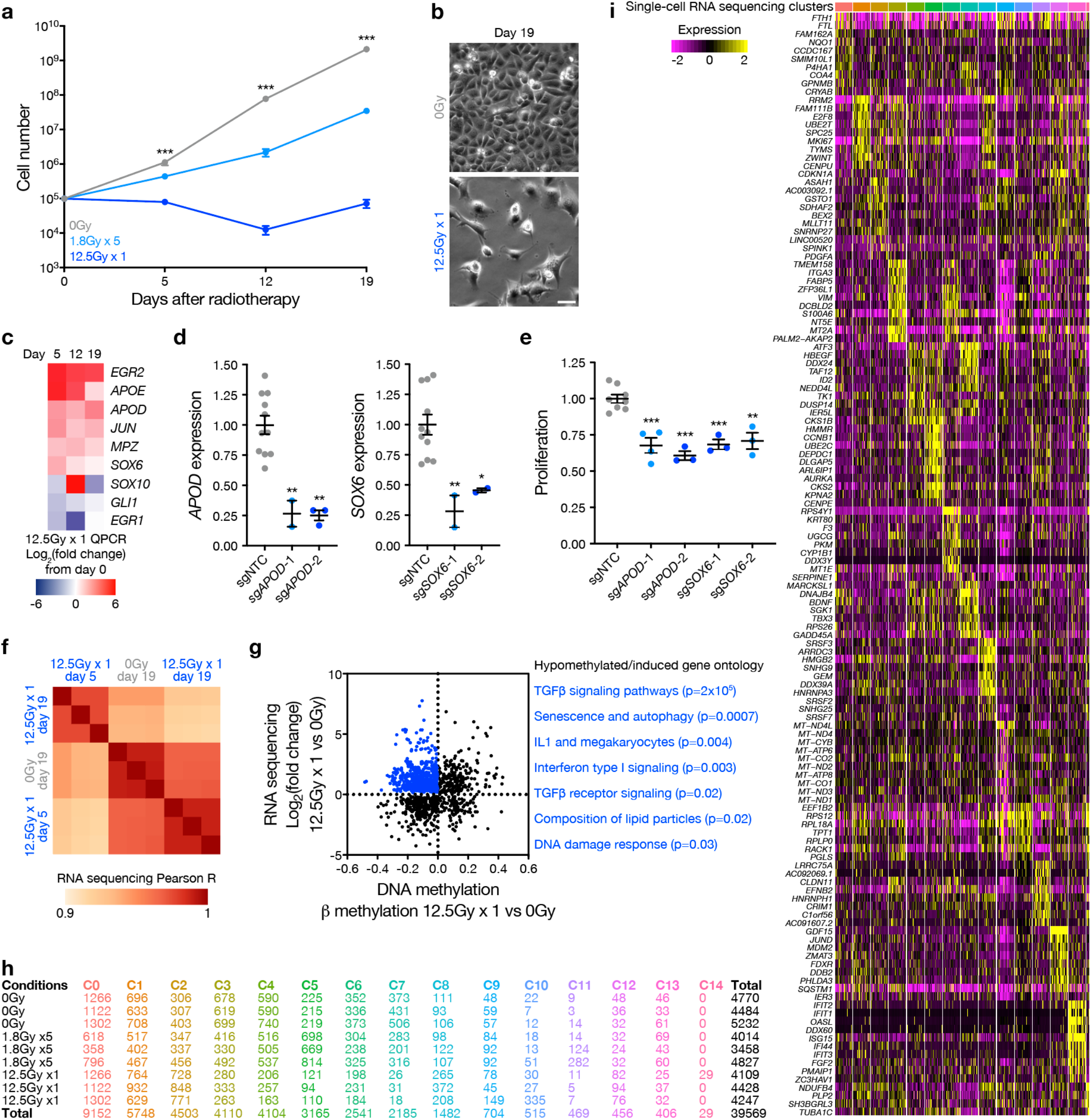
Radiotherapy epigenetically reprograms schwannoma cells to multiple cellular states, expressing immune and inflammatory genes that regulate schwannoma cell growth. **a**, Quantification of HEI-193 schwannoma cell proliferation over time after treatment with 0Gy, 1.8Gy x 5, or 12.5Gy x1. **b**, Light microscopy of HEI-193 cells treated with either 0Gy or 12.5Gy x1 of radiotherapy. Scale bar, 10μm. **c**, Heatmap of log_2_ fold changes of QPCR gene expression between day 0 and day 5, 12 or 19 following 12.5Gy x1 of radiotherapy treatment of HEI-193 cells. Heatmap values represent averages of 3 QPCR replicates per condition. **d**, CRISPRi suppression *APOD* or *SOX6* in HEI-193 cells compared to non-targeted control sgRNAs (sgNTC). **e**, Quantification of HEI-193 cell proliferation after CRISPRi repression of *APOD* or *SOX6* using 2 independent sgRNAs per gene as in **d**, compared to sgNTC. Lines represent means and error bars represent standard error of means (Student’s t tests, *p:≤0.05, **p:≤0.01, ***p:≤0.0001). **f**, Heatmap of pairwise Pearson correlation coefficients between RNA sequencing of triplicate HEI-193 cultures after treatment with either 0Gy or 12.5Gy x 1 radiotherapy, 5- or 19-days following radiation treatment. **g**, Comparison of RNA sequencing and DNA methylation profiling of HEI-193 cells 19 days after 12.5Gy x 1 of radiotherapy compared to 0Gy control treatment. Significant gene ontology terms for enriched and hypomethylated genes are shown (blue, n=3 replicate cultures per condition). **h**, Distribution of HEI-193 single-cell RNA sequencing transcriptomes (Fig. 2c). **i**, Expression heatmap of schwannoma cell state marker genes from single-cell RNA sequencing of triplicate HEI-193 cultures following 0Gy, 1.8Gy x 5, or 12.5Gy x 1 radiotherapy. Columns represent transcriptomes from single cells downsampled to 200 cells per cluster.

**Extended Data Fig. 8.**
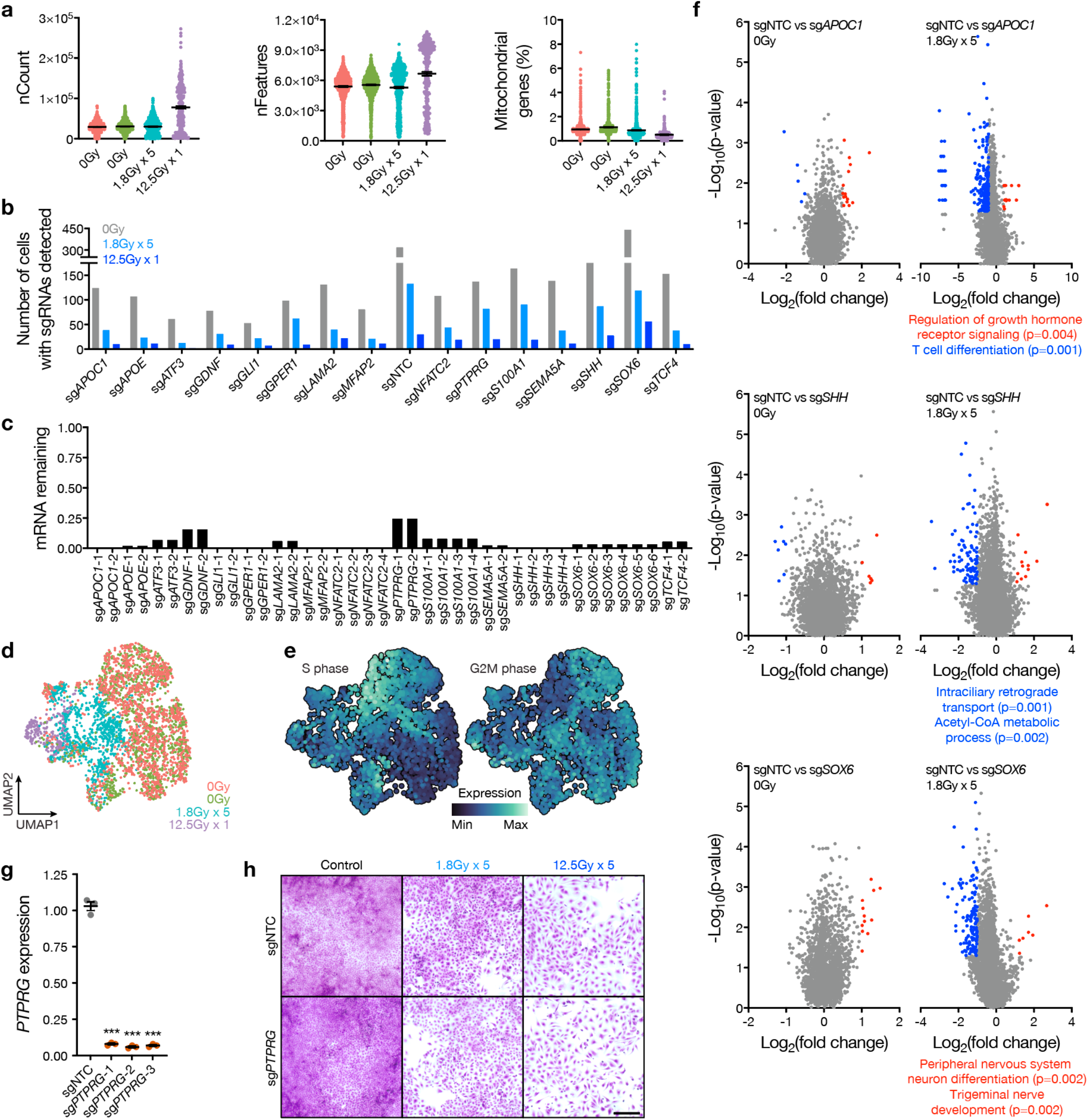
Perturb-seq of schwannoma marker genes reveals regulators of cell states. **a**, Distributions of UMI counts (left), number of unique gene expression features (middle), and percentage of UMIs mapping to mitochondria gene (right) from Perturb-seq in HEI-193 cells. **b**, sgRNA coverage for each targeted gene in 0Gy, 1.8Gy x 5, or 12.5Gy x 1 radiotherapy conditions. n=2-6 sgRNAs per gene targeted. **c**, Mean knockdown of targeted genes in cells pseudobulked by each sgRNA. Only sgRNAs with confirmed repression of greater than 75% were retained for analysis. **d**, UMAP of 3546 HEI-193 cells passing quality thresholds for Perturb-seq. **e**, Feature plot of S phase or G2M phase genes in Perturb-seq UMAP from **d. f**, Volcano plots for differential gene expression analysis using MAST for *APOC1, SHH*, or *SOX6* perturbations compared to sgNTC in 0Gy (left) versus 1.8Gy x 5 conditions (right). Significant positive (red) or negative (blue) gene expression changes are colored (p<0.05, |log_2_(fold change)|>1), corresponding to gene ontology terms. **g**, CRISPRi suppression of *PTPRG* in HEI-193 cells compared to non-targeted control sgRNAs (sgNTC). **h**, Representative crystal violet staining of HEI-193 cells following CRISPRi suppression of *PTPRG* compared to sgNTC after treatment with 0Gy, 1.8Gy x 5, or 12.5Gy x 1 of radiotherapy. Scale bar, 100μm. Lines represent means and error bars represent standard error of means (Student’s t tests, ***p:≤0.0001).

**Extended Data Fig. 9.**
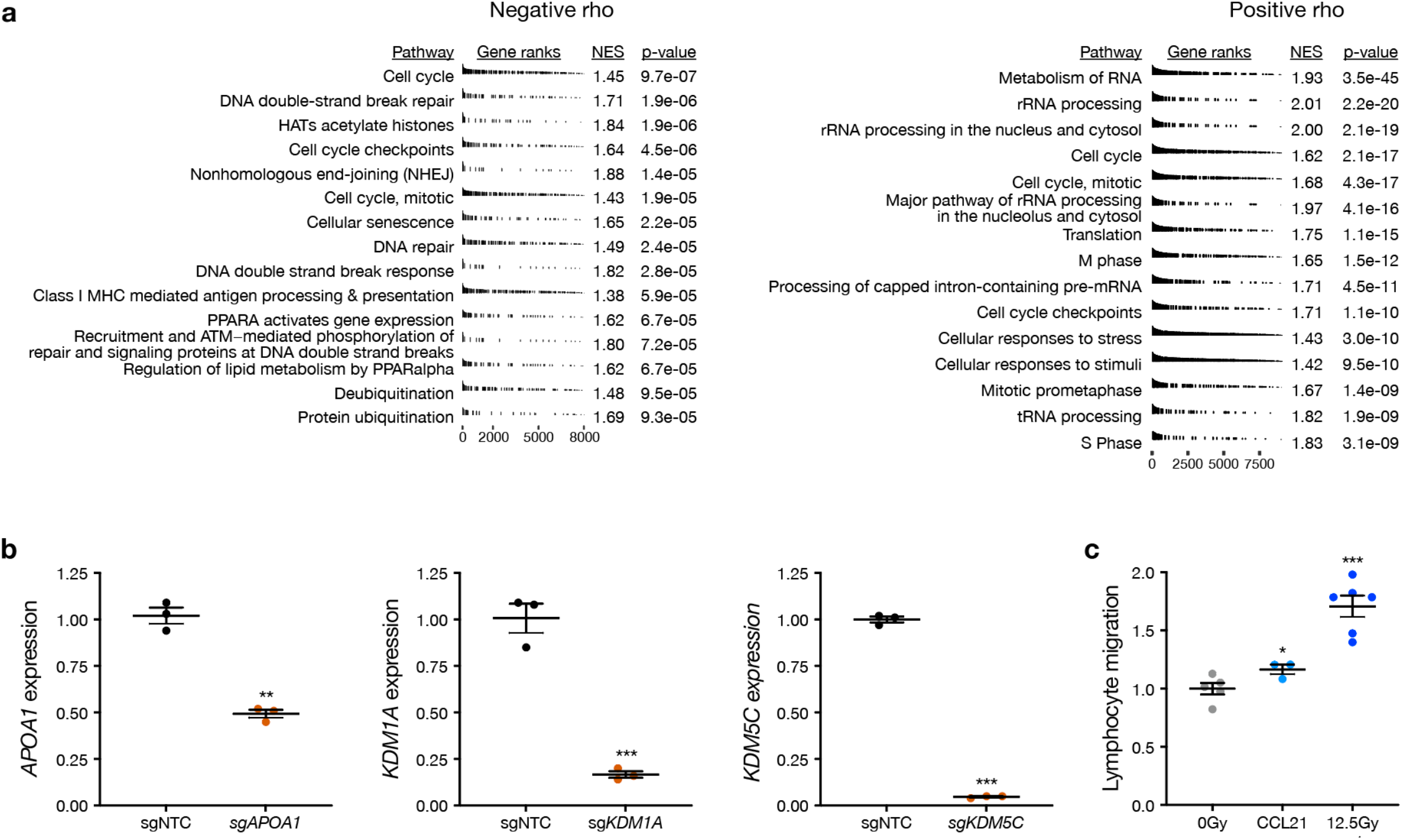
Genome-wide CRISPRi screens reveal regulators of schwannoma cell radiotherapy responses. **a**, Gene set enrichment analysis of ranked gene targets exhibiting negative rho (left, radiotherapy sensitivity) or positive rho (right, radiotherapy resistance) following CRISPRi gene suppression. Gene ontology terms derived from Reactome. NES, normalized enrichment score. **b**, CRISPRi suppression of *APOA1, KDM1A*, or *KDM5C* in HEI-193 cells. **c**, Transwell primary human peripheral blood lymphocyte migration assays using conditioned media from HEI-193 cells *±* radiotherapy or with recombinant CCL21 as a chemoattractant. Lines represent means and error bars represent standard error of means (Student’s t tests, *p:≤0.05, **p:≤0.01, ***p:≤0.0001).

**Extended Data Fig. 10.**
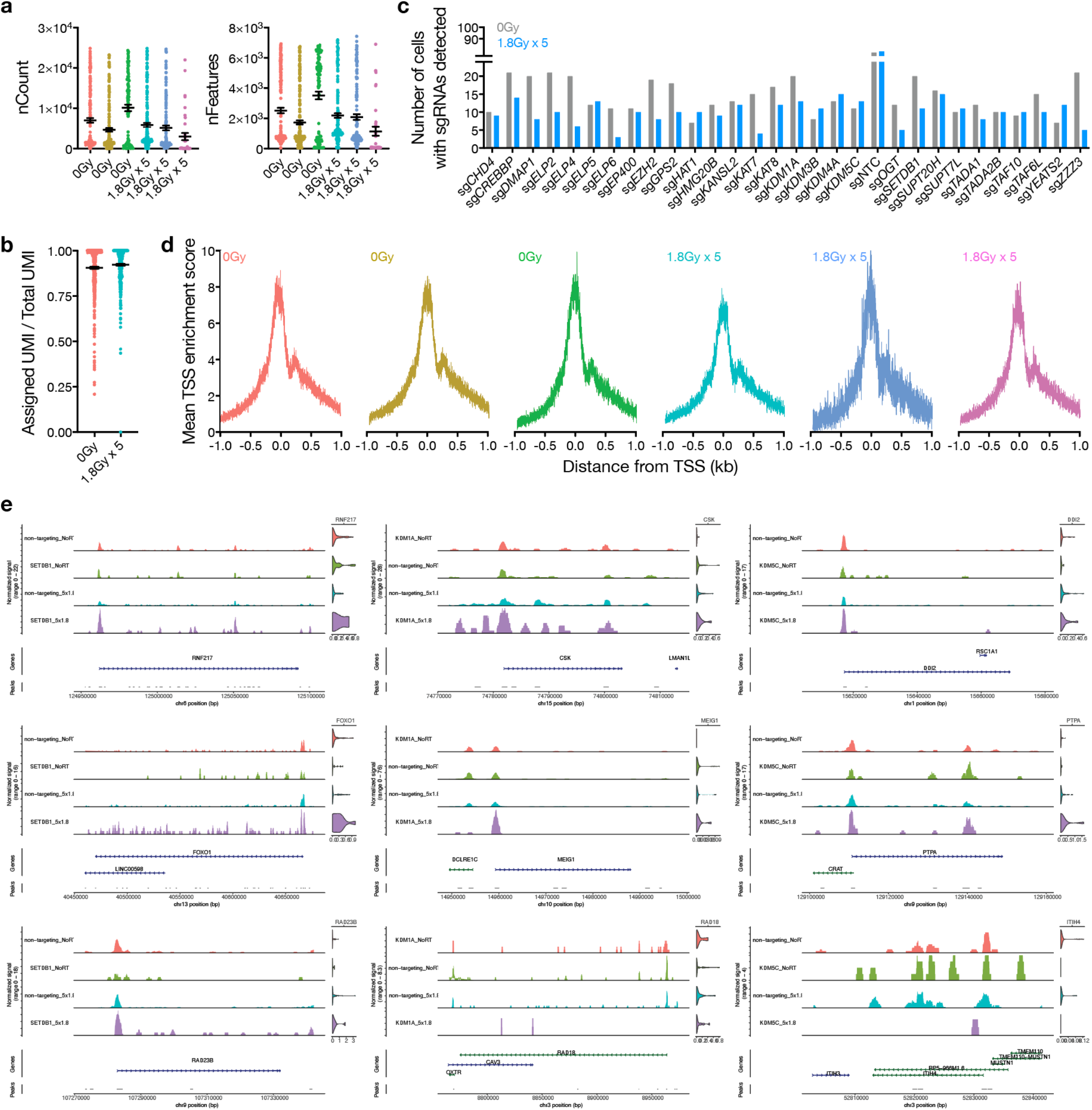
snARC-seq quality metrics for simultaneous profiling of chromatin accessibility and gene expression in the context of therapeutic and CRISPRi perturbations. **a**, Distributions of UMI count (left) and number of unique gene expression features (right) from snARC-seq in HEI-193 cells. **b**, Distributions of UMIs mapping to sgRNAs as a proportion of all sgRNA UMI counts in individual HEI-193 cells. Lines represent means and error bars represent standard error of means. **c**, sgRNA coverage for each targeted gene in 0Gy or 1.8Gy x 5 radiotherapy conditions. **d**, Transcription start site (TSS) enrichment profile plots for snARC-seq ATAC chromatin accessibility profiles, demonstrating characteristic nucleosome-free regions relative to the TSS in each condition. **e**, Pseudobulked ATAC chromatin accessibility profiles coupled with single-nuclei RNA expression distributions at example genomic loci of genes that were activated with *SETDB1* perturbation (left), activated or repressed with *KDM1A* perturbation (middle), or activated or repressed with *KDM5C* perturbation (right), in the presence or absence of radiotherapy (1.8Gy x 5).

**Supplementary Fig. 1.**
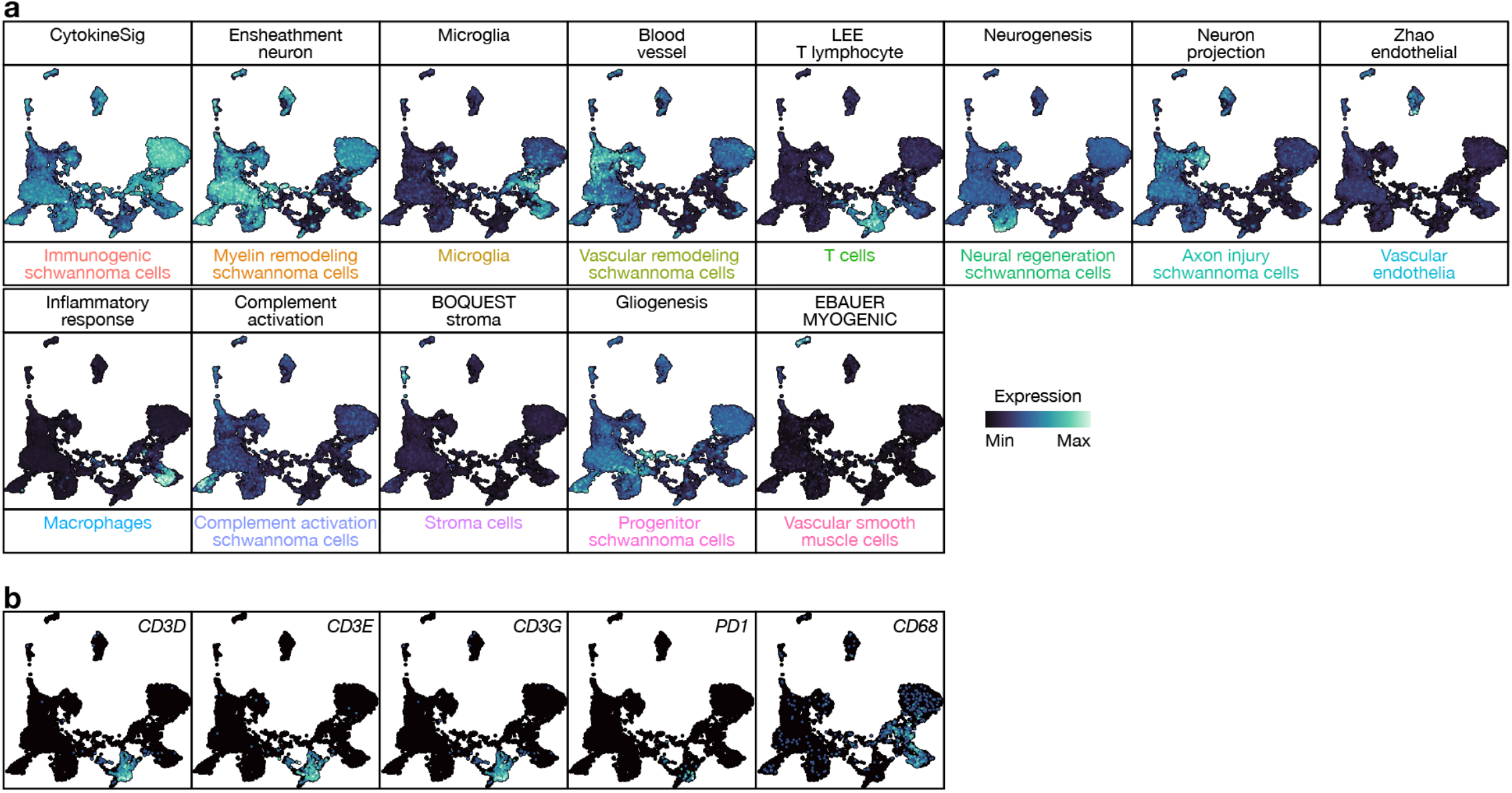
Integrated schwannoma single-nuclei and single-cell RNA sequencing feature plots. **a**, Normalized gene expression feature plots of signatures derived from Molecular Signatures Database^80^ gene sets intersecting with top cluster markers from integrated single-nuclei and single-cell RNA sequencing of human schwannomas supporting schwannoma cell type or microenvironment cell type definitions. **b**, Gene expression feature plots of lymphoid or myeloid marker genes supporting immune cell type definitions. Related to Fig. 1b.

**Supplementary Fig. 2.**
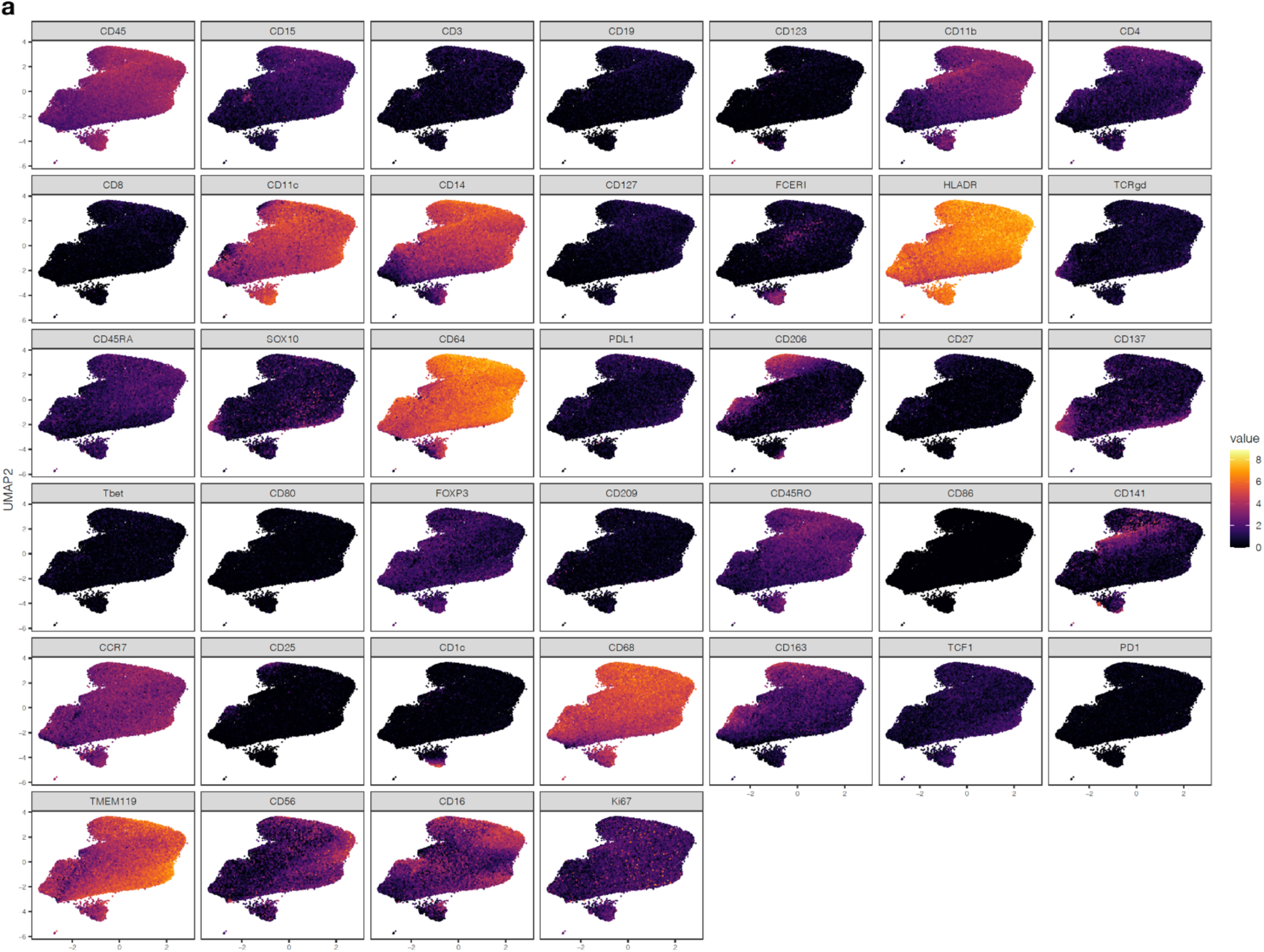
CyTOF myeloid cell feature plots. Related to Fig. 1f, g. Legend shows asinh intensity.

**Supplementary Fig. 3.**
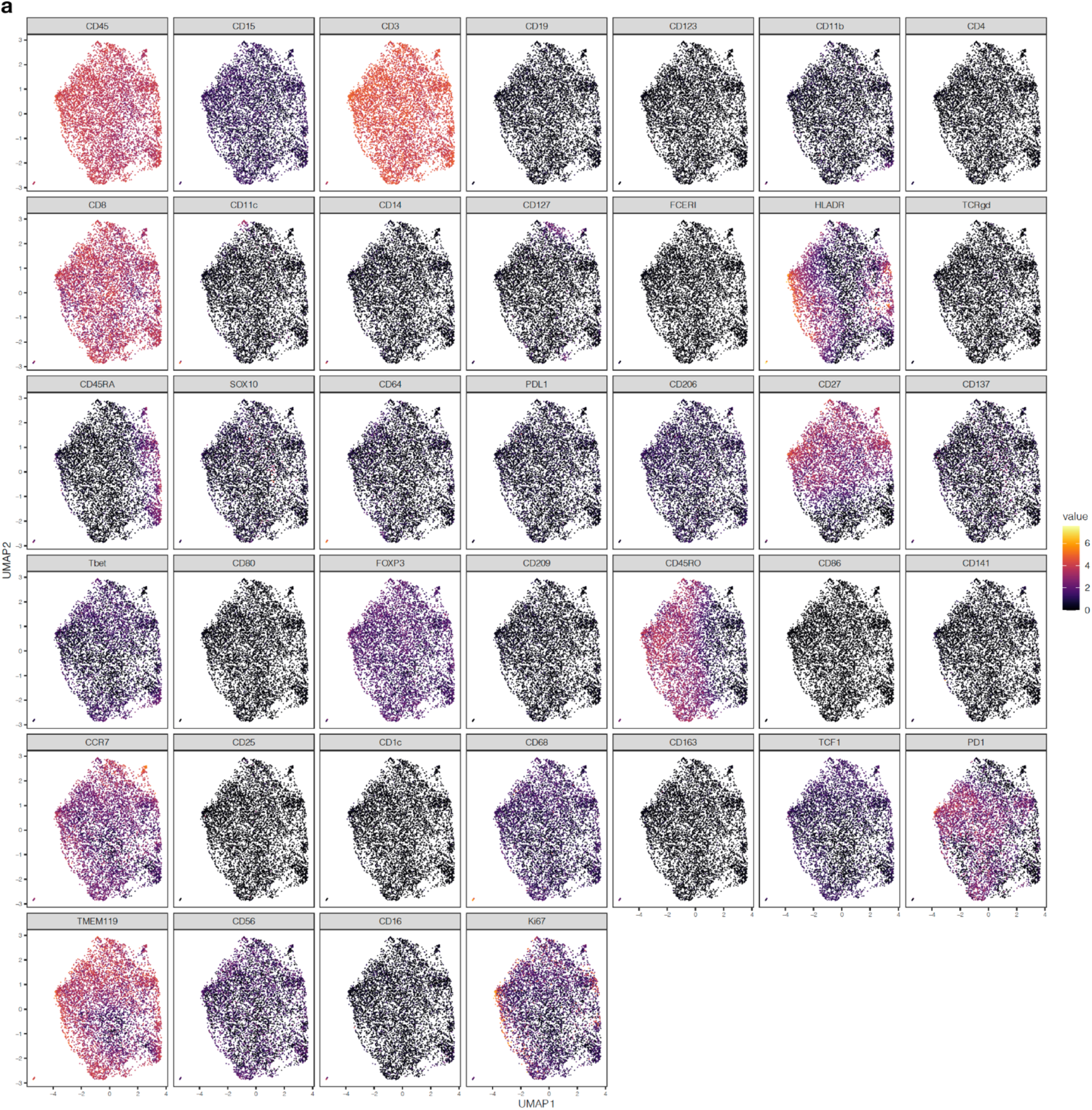
CyTOF CD8 T cell feature plots. Related to Fig. 1f, g. Legend shows asinh intensity.

**Supplementary Fig. 4.**
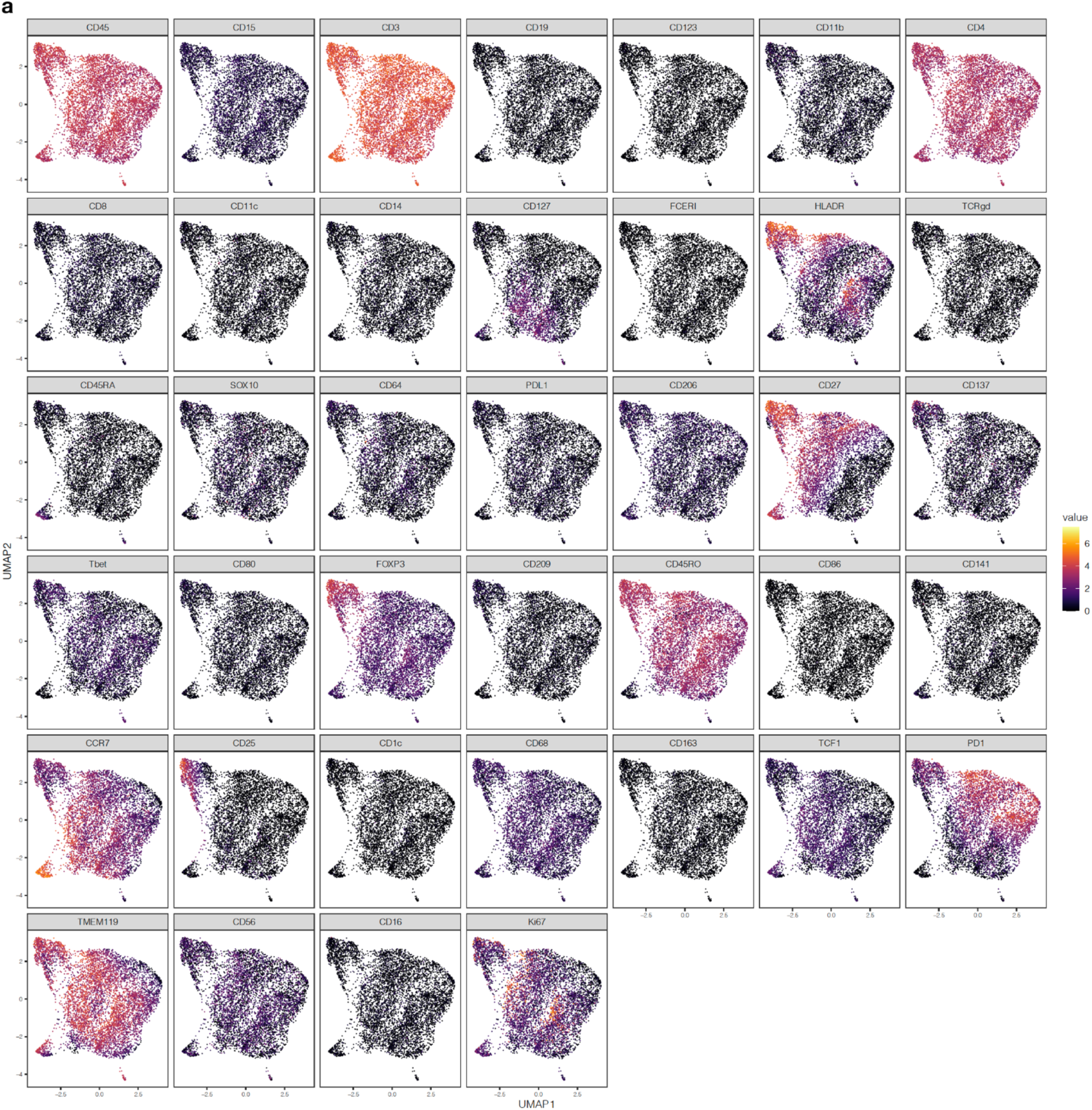
CyTOF feature plots used to define CD4 T cells. Related to Fig. 1f, g. No significant differences in CD4 T cell enrichment were identified between NCS and IES. Legend shows asinh intensity.

**Supplementary Fig. 5.**
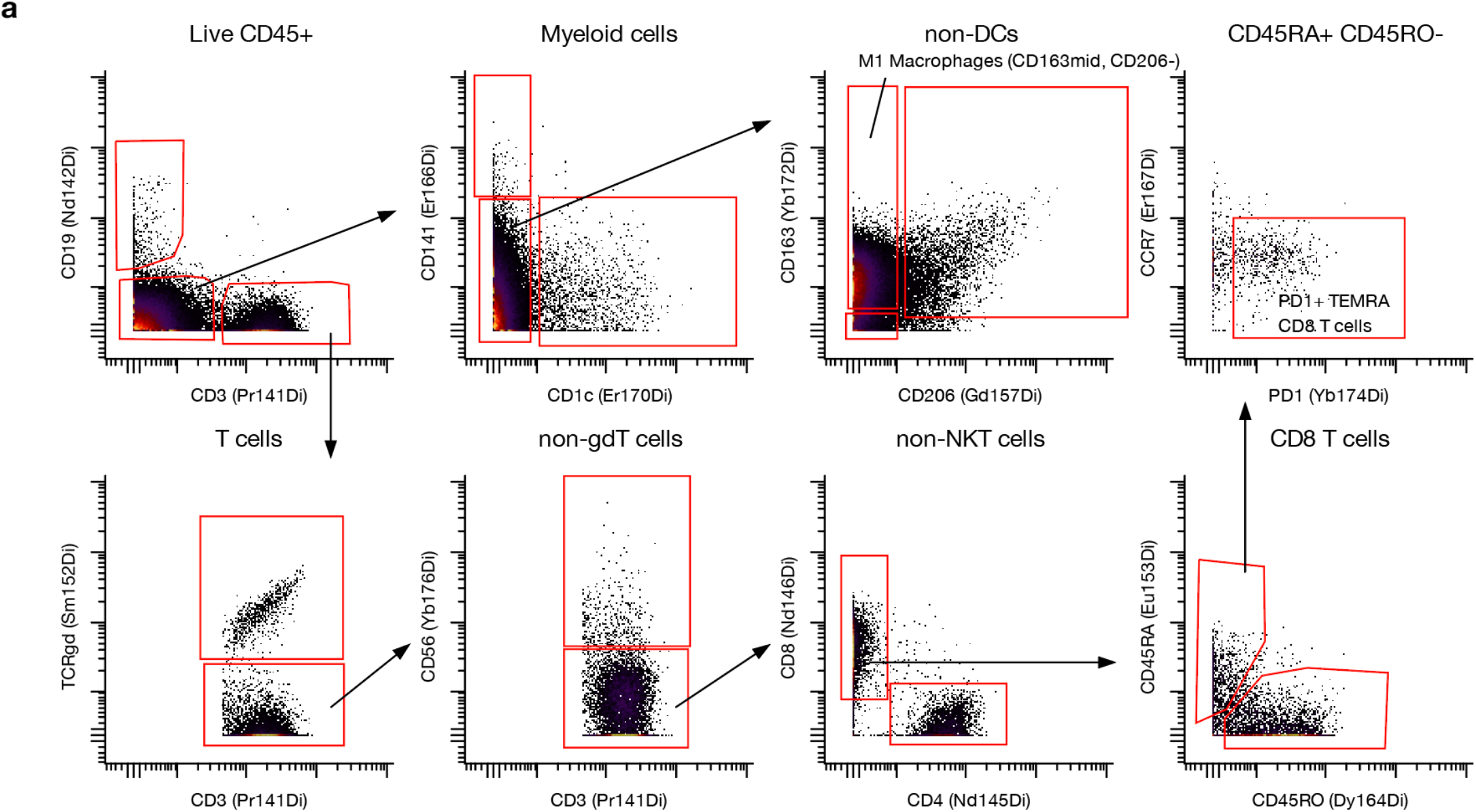
CyTOF gating workflow used to define schwannoma immune cell populations such as myeloid cells and CD8 T cells. Related to Fig. 1f, g.

**Supplementary Fig. 6.**
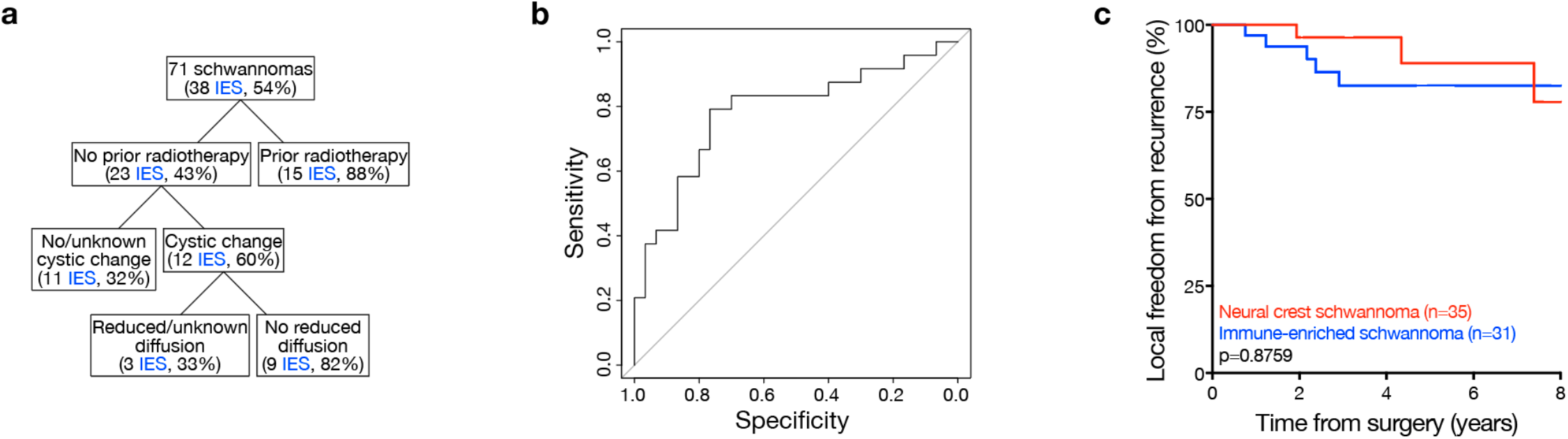
Analysis of clinical features distinguishing schwannoma molecular groups. **a**, Recursive partitioning analysis of clinical and magnetic resonance imaging features in patients with IES versus NCS (n=71). **b**, Receiver operating characteristic curve for logistic regression modeling to predict schwannoma molecular group based on non-invasive clinical and magnetic resonance imaging features (area under the curve 0.79). c, Kaplan-Meier curves for local freedom from recurrence for NCS versus IES demonstrating no significant difference in tumor control between molecular groups (log-rank test).

**Supplementary Fig. 7.**
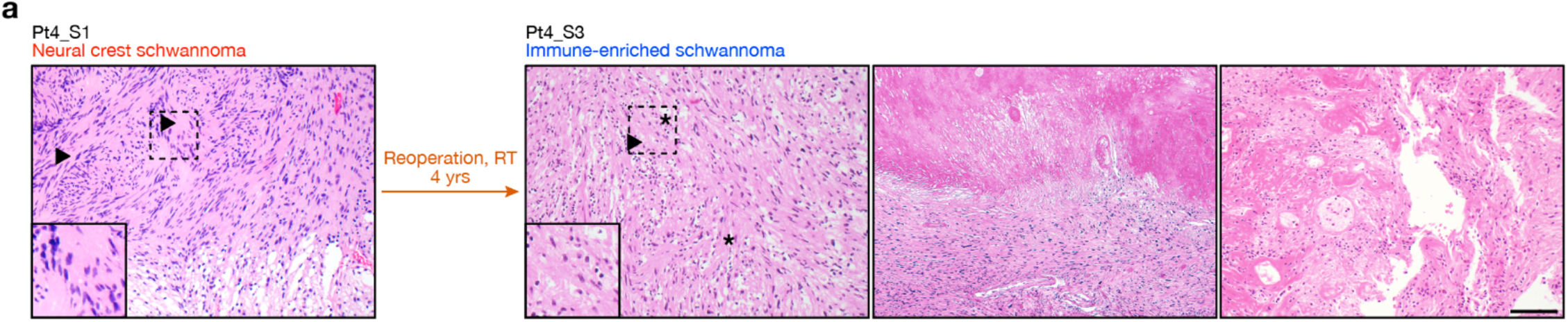
Radiotherapy durably recruits immune cells to the schwannoma microenvironment. **a**, H&E-stained sections of a patient-matched primary NCS and recurrent IES, showing typical histology with biphasic histology and Verocay bodies (left arrows) that was replaced by abundant lymphocytes (right arrows) with foamy and hemosiderin-filled macrophages (asterisks), coagulative necrosis (second from right), and hyalinized vessels (far right). RT, radiotherapy. Scale bar, 100μm.

**Supplementary Fig. 8.**
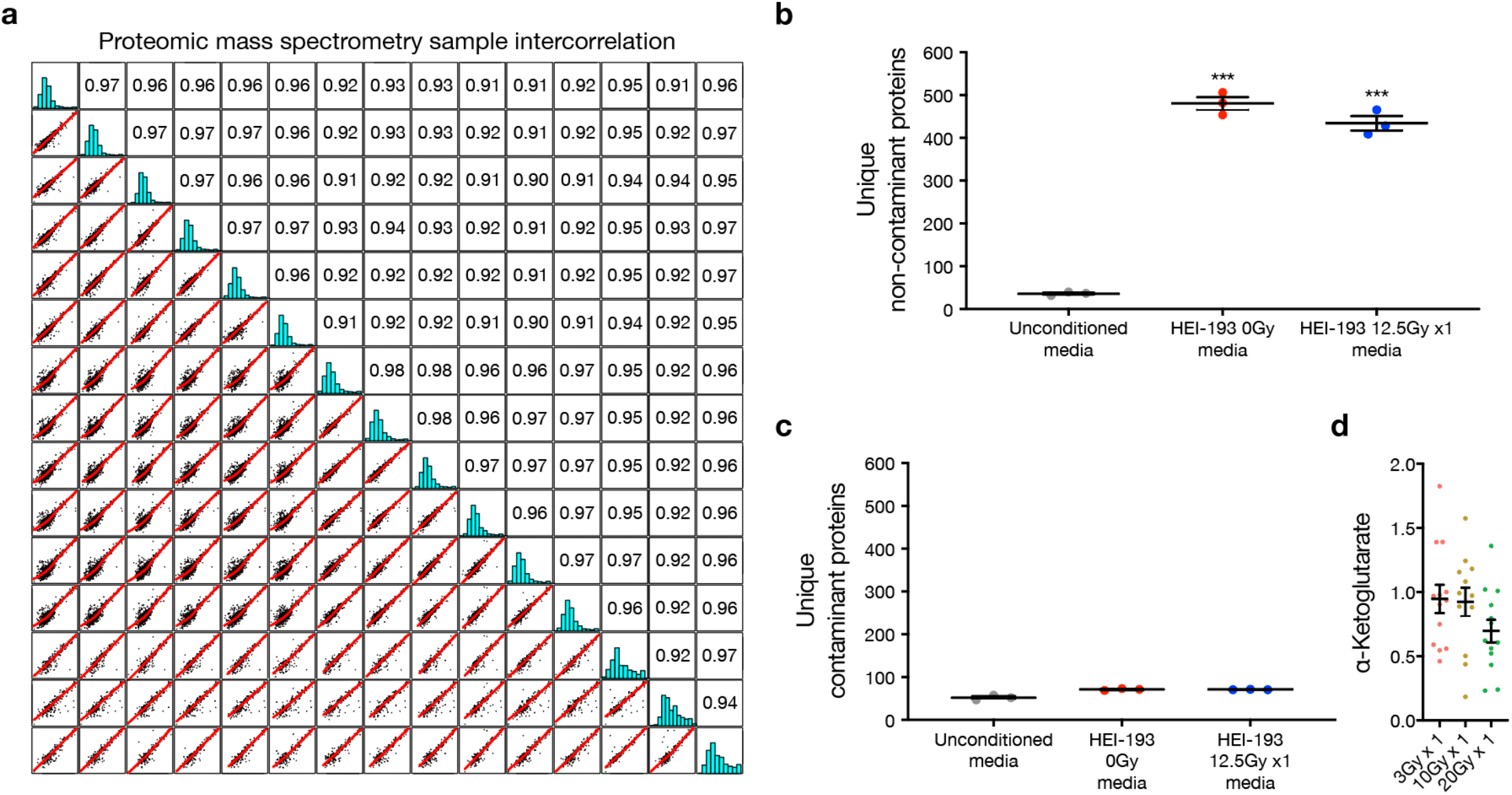
Proteomic and targeted metabolic mass spectrometry of schwannoma cells. **a**, Proteomic mass spectrometry Pearson correlation matrix from HSC and HEI-193 media demonstrating high correlation across samples. **b** and **c**, Unique non-contaminant or contaminant protein counts from HEI-193 media mass spectrometry (Student’s t tests, ***p*≤*0.0001). **d**, Metabolite mass spectrometry of primary patient-derived human schwannoma cells (n=7) after treatment with 3Gy x 1, 10Gy x 1, or 20Gy x 1 of radiotherapy validated suppression of α-Ketoglutarate with ionizing radiation (Fig. 3i). Fold changes normalized to 0Gy treatment for each primary cell culture (ANOVA, p≤0.05). Lines represent means and error bars represent standard error of means.

## Notes

### Competing Interest Statement

The authors have declared no competing interest.

